# The Phylogenomics of CRISPR-Cas System in *Salmonella*: an evolutionary race with housekeeping genes

**DOI:** 10.1101/2020.07.08.193417

**Authors:** Simran Krishnakant Kushwaha, Chandrajit Lahiri, Bahaa Abdella, Sandhya Amol Marathe

## Abstract

*Salmonellae* display intricate evolutionary patterns comprising over 2500 serovars having diverse pathogenic profiles. The acquisition/exchange of various virulence factors influence the evolutionary framework. To gain insights into evolution of *Salmonella* as a pathogen in association with the CRISPR-Cas genes we performed phylogenetic surveillance across strains of 22 *Salmonella* serovars. The strains assorted into two main clades, pertaining to the differences in their CRISPR1-leader and *cas* operon. Considering *Salmonella enterica* subsp. *enterica* serovar Typhimurium and serovar Typhi as signature serovars, we classified the clades as CRISPR1-STM/*cas*-STM and CRISPR1-STY/*cas*-STY, respectively. Serovars of the two clades displayed better relatedness, concerning CRISPR-1 leader and *cas* operon, across genera than between themselves. This signifies the acquisition of CRISPR1/Cas region a horizontal gene transfer event owing to the presence of mobile genetic elements flanking CRISPR1 array. The CRISPR2 tree does not show such relation. Spacer mapping of the two CRISPR arrays suggests the construct to be canonical, with only 8.8% spacer conservation among the serovars. As opposed to broad-host-range serovars, the host-specific serovars harbor fewer spacers. All typhoidal serovars have CRISPR1-STY/*cas*-STY system. Comparison of CRISPR and *cas* phenograms with that of multilocus sequence typing (MLST) suggests differential evolution of CRISPR/Cas system implying supplementary roles beyond immunity.

## 1. INTRODUCTION

Genus *Salmonella* is classified into two species, *Salmonella enterica* and *Salmonella bongori*. *S. enterica* further evolved into six subspecies (subsp.) *enterica*, *salamae*, *arizonae*, *diarizonae*, *houtenae* and *indica*^1^. Serovars of *S. enterica* subsp. *enterica* have diverse host range ranging from broad-host-range, host-adapted to host-specific^2^ pertinent to within-host evolution^3^.

Before divergence, *S. bongori* and *S. enterica* acquired *Salmonella* pathogenicity island 1 (SPI-1)^4^ and later *S. enterica* laterally acquired SPI-2 (required for the survival in deep tissues and macrophages) thereby, enhancing its virulence potential^4^. As per the adopt-adapt model of bacterial speciation^5^, the adopted lateral gene(s) divert the evolution path promoting bacterial adaptation consequently increasing its fitness^6^. Over time, *S. bongori* and *S. enterica* horizontally acquired multiple virulence factors progressively enhancing their pathogenicity^3^.

The presence of the Clustered Regularly Interspaced Short Palindromic Repeats (CRISPR) and a cognate set of CRISPR-associated (*cas*) genes, has recently been related to the bacterial virulence potential^7^. In *E. coli*, CRISPR units are negatively correlated with pathogenic potential where, the reduction in CRISPR activity is proposed to promote lateral gene transfer favoring its evolution^8^. Conversely, some reports demonstrate a positive correlation between the CRISPR and pathogenicity owing to virulence genes regulation^9^. *S. enterica* possesses type I–E CRISPR system with two CRISPR arrays, CRISPR1 and CRISPR2^10^, separated by ~16 kb^11^. Many proto-spacers of this were traced on the chromosome instead of its general target - phages and plasmids^12^. This suggests a potential role of the CRISPR-Cas system in endogenous (virulence) gene regulation possibly regulating the pathogenesis of *Salmonella*^13^. To test the association of CRISPR-Cas system and pathogenicity, we studied the evolutionary dynamics of CRISPR-Cas system across 47 strains of *Salmonella* species. The evolution was studied in the light of three essential attributes of CRISPR-Cas components the leader, the array, and the *cas* operon. We created a graphic map of the spacers of 131 strains across 22 serovars belonging to two species of *Salmonella*, comprehending the structural composition and its configuration. Next, the strains were assorted into two groups with respect to the CRISPR1 leader, spacer composition, and *cas* gene composition and similarity. This divergence was analyzed in comparison to multi-locus sequence typing (MLST) based on the seven core genes. Our study elucidates the hidden aspects and develops a strong correlation between the CRISPR-Cas system and the host specificity of various serovars of *Salmonella,* demonstrating that this system could have potential involvement in the virulence and pathogenicity.

## 2. RESULTS

### 2.1 Diversity of the CRISPR arrays in *Salmonella*

We extracted all possible CRISPR1 and 2 arrays in correct orientation for 131 strains (Table S1, supplementary methodology). *S. bongori* and *S. enterica* subsp. *enterica* contained both CRISPR arrays while subsp. *arizonae* and subsp. *diarizonae,* had only one CRISPR array. One strain (str. NCTC10047), out of the six strains examined of subspecies *arizonae*, and all the eight strains of subspecies *diarizonae* had an intact CRISPR array.

We mapped the spacer sequences (Fig. S1) of all 131 strains illustrating the blueprint of spacer conservation among the strains within and across serovars. The acquisition of spacers is in a precise fashion with conserved configuration for a particular serovar while the differences lie in the presence or absence of spacer(s). The spacers of serovars Enteritidis, Heidelberg, and Typhi are highly conserved among all its strains, whereas the serovars Typhimurium, Newport, Anatum, Montevideo and Tennessee had significant variability in the spacer composition of both the CRISPR arrays (Fig. 1 and Fig. S1a). Among all strains, we identified 437 and 327 unique spacers in CRISPR1 and CRISPR2 arrays, respectively. The average abundance of spacers for CRISPR1 and CRISPR2 is 15.3 and 12.6, respectively (Table S2). CRISPR1 array of serovar Tennessee str. ATCC 10722 (63 spacers) and CRISPR2 array of serovar Typhimurium str. USDA-ARS-USMARC-1880 (35 spacers) are the largest (Fig. S1). CRISRP1 array of serovar Anatum, Dublin, Gallinarum, Pullorum and, Gallinarum/Pullorum (two spacers), and CRISPR2 array of serovars Sendai and Typhi (one spacer) are the shortest (Fig. S1). We observed duplication and triplication of spacer in some serovars (Fig. S1a-S1b).

**FIGURE 1.**
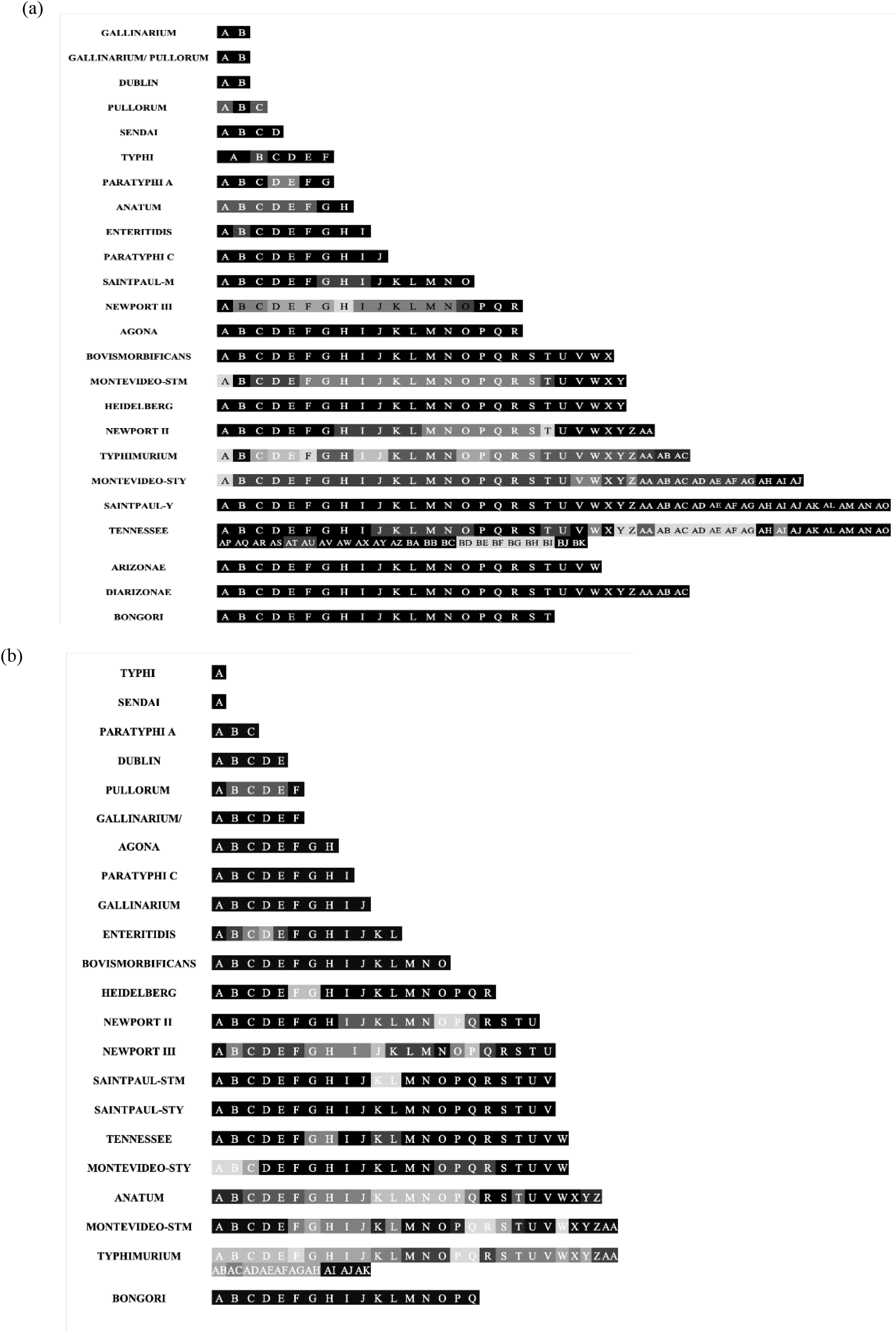
Graphic map of spacer conservation in CRISPR1 (**a**) and CRISPR2 (**b**) array for *Salmonella* serovars. The shades of grey represent the conservation percentage of a given spacer in all the strains of the respective serovar where, the darker box indicate the presence of spacer in most of the strains (black: 100%) while, the lighter box indicate the presence of spacer in a few strains.

Strikingly, the analysis of the CRISPR arrays in serovars Montevideo and Saintpaul separated the respective strains into two groups with two distinct sets of unique and conserved spacers (Table S1). For serovar Montevideo, the two groups comprised eight and nine strains. However, CRISPR arrays of all the analyzed strains of serovar Saintpaul except strain SARA26 (an outlier) had similar spacer composition. These results suggest polyphyletic origin of serovars Montevideo and Saintpaul, similar to that reported for serovar Newport^14^. Notably, the broad-host-range serovars have multiple spacers, while the host-specific serovars have very few spacers (Fig. 1).

A bird’s eye view on direct repeat (DR) sequence of the CRISPR arrays confirmed the sequence to be 29 bp long, partially palindromic^12^ and the last DR is degenerate^15^(Fig. S2b, S2d & S2f). The DR sequence is conserved within respective array across all the serovars except for the presence of few SNPs (Fig. S2a, S2c & S2e). The degeneration of last DR is of varying degrees, primarily witnessed near the 3′ end, while the 5′ end substantially conserved. Considerable difference in the last DR of both the arrays (Fig. S2) suggests existence of heterogeneous termination mechanism of pre-crRNA processing

### 2.2 Phylogeny and classification of the CRISPR arrays

#### 2.2.1 Evolutionary studies of the CRISPR loci

To understand the evolutionary pattern of *Salmonella* serovars concerning the CRISPR loci, we generated phylogenetic trees inclusive of the array and leader sequence (Fig. 2) for 47 shortlisted set of strains (Table S1). These robust phylograms signify predominant divergence of the CRISPR loci. The topology has been observed in most of the clades and sub-clades, as evidenced by their high level of confidence from either the bootstrap values or the aLRT (approximate likelihood ratio test) scores.

**FIGURE 2.**
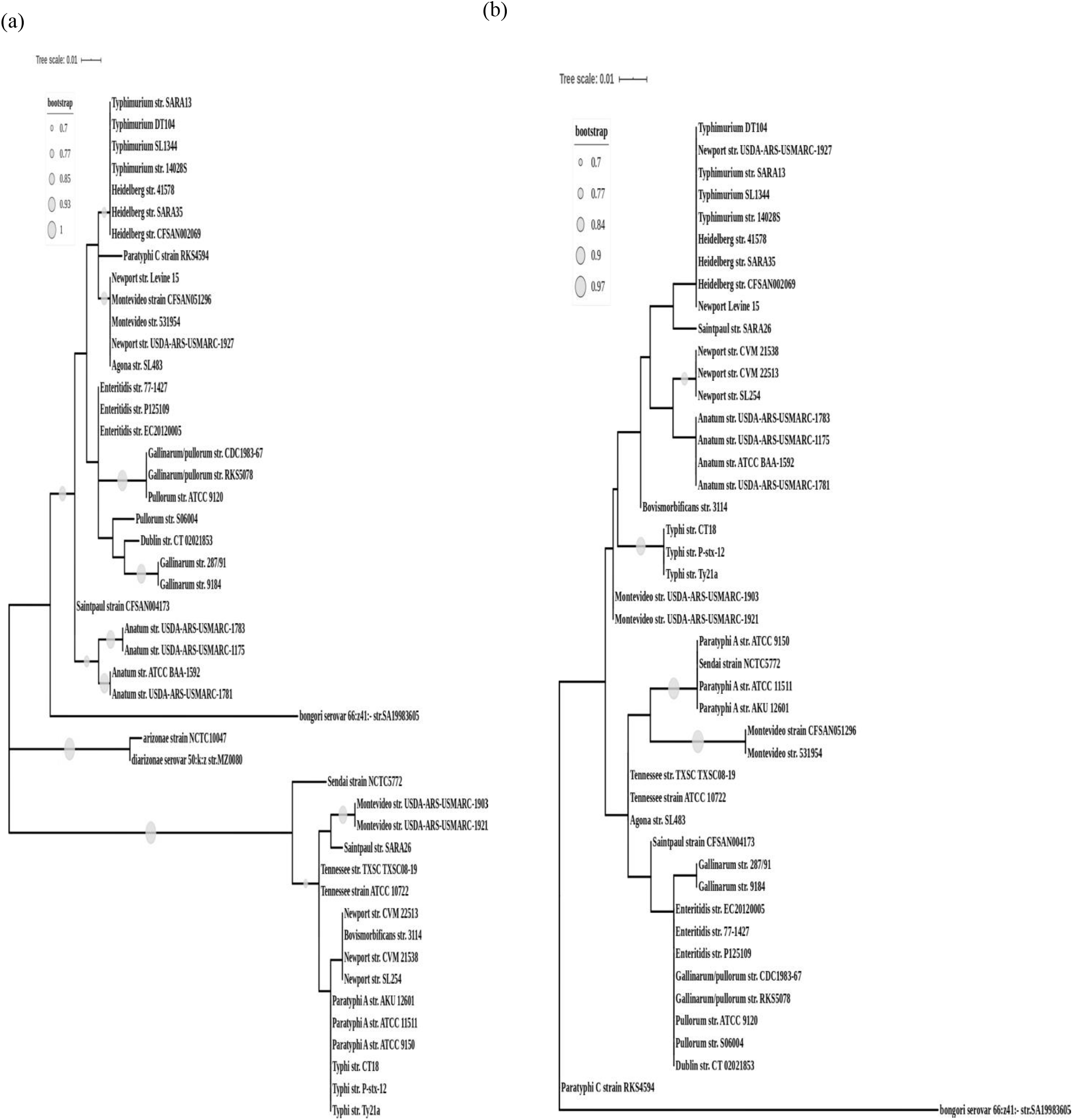
The phylogeny of CRISPR leader+array, CRISPR1 (**a**), and CRISPR2 (**b**) across *Salmonella* serovars. The entire CRISPR arrays and their leader sequences were aligned using Muscle, and the phylogenetic trees were constructed by ML. The trees were visualised by iTOL.

The CRISPR1 tree had two distinct clades with serovars Typhimurium and Typhi as signature serovars (Fig.2). Thus, we classified the corresponding CRISPR loci as CRISPR1-STM and CRISPR1-STY, respectively. The strains of serovars Saintpaul and Montevideo harboring these loci were defined as Saintpaul-STM/ Montevideo-STM, and Saintpaul-STY/ Montevideo-STY, respectively.

In CRISPR 2 phenogram (Fig. 2), *S. bongori* emerged as an outgroup for the entire tree, and serovar Paratyphi C seems to have evolved distinctly from other serovars of *S. enterica* subsp. *enterica*. The CRISPR2 tree topology and sub-lineages were very distinct from that of the CRISPR1 tree with serovars forming polyphyletic clades. For example, serovar Saintpaul-STY grouped with serovars Typhimurium, Newport III, Heidelberg; Sendai, Paratyphi A grouped with Montevideo-STM; and Newport II with Anatum. The clustering was distinct from that obtained in the CRISPR1 tree. This can be authenticated by the aLRT scores evaluating that each branch under review provides a significant likelihood gain (Fig. S3 & S4) suggesting different evolutionary relationships of both the CRISPR loci.

#### 2.2.2. Comparison of the spacer sequences in the light of CRISPR phylogeny

To check if the difference in the CRISPR1 and CRISPR2 trees is due to the differential spacer conservation, the inter-serovar spacer content was analyzed (query cover >95%). Only ~8-9.5% of unique spacers (36/437: CRISPR1 and 31/327: CRISPR2) were shared by two or more serovars (Fig. S1C). Some serovars in CRISPR1-STM clade have common spacers justifying their clustering. For example, serovars Typhimurium, Heidelberg, Newport III, and Paratyphi C; and serovars Anatum and Saintpaul-STM share their anchor spacer. Entire spacer set of serovars Gallinarum, Pullorum, and Gallinarum/Pullorum are present in serovar Enteritidis (Fig. S1c). Similarly, few serovars of CRISPR1-STY clade share some spacers, e.g., serovars Paratyphi A and Sendai; and serovars Bovismorbificans and Newport II; share spacers, but these serovars do not cluster as expected.

For the CRISPR2 tree, atypically, the first spacer of serovars Enteritidis, Gallinarum, Pullorum, and Gallinarum/Pullorum; and serovar Paratyphi A and serovar Sendai is conserved (Fig. S1d). Serovars Paratyphi A, Saintpaul-STY, Montevideo-STY, and Newport II have one spacer in common, which is the last spacer in all the serovars except Newport II. Serovars Typhi, Enteritidis, Paratyphi C, Gallinarum, Pullorum and, Gallinarum/Pullorum share their anchor spacer. However, the serovars with identical anchor spacer belong to different clusters.

In the CRISPR2 locus tree and CRISPR1-STY clade, the serovars are not always clubbed as per the spacer similarity.

#### 2.2.3 Categorization of the leader sequence in the light of CRISPR phylogeny

Multiple sequence alignment of respective CRISPR sequences across the serovars revealed a significant match in the DR and the leader sequence. The CRISPR1 DR is conserved across the strains of CRISPR1-STM, and CRISPR1-STY clades with an SNP in a few strains of serovar Anatum (Fig. S2) thereby indicating major influence of leader on the topology of the CRISPR1 tree. To confirm this, we constructed phylogenetic trees for CRISPR1 and CRISPR2 leaders (Fig. 3). The topology of the CRISPR2 leader tree more or less coincided with the CRISPR2 tree (Fig. 2 and 3; supplementary figure S4 and S7). The clustering pattern of serovars in the CRISPR1 tree and CRISPR1 leader tree is similar except higher substitutions and enhanced branching in CRISPR1 tree (Fig. 2 and 3; supplementary figure S3 and S6). This indicates that the major force for segregation of CRISPR1-STM and CRISPR1-STY clades is the variation in the leader sequence, and the formation of sub-clades/sub-clusters is a result of shared spacers among the serovars.

**FIGURE 3.**
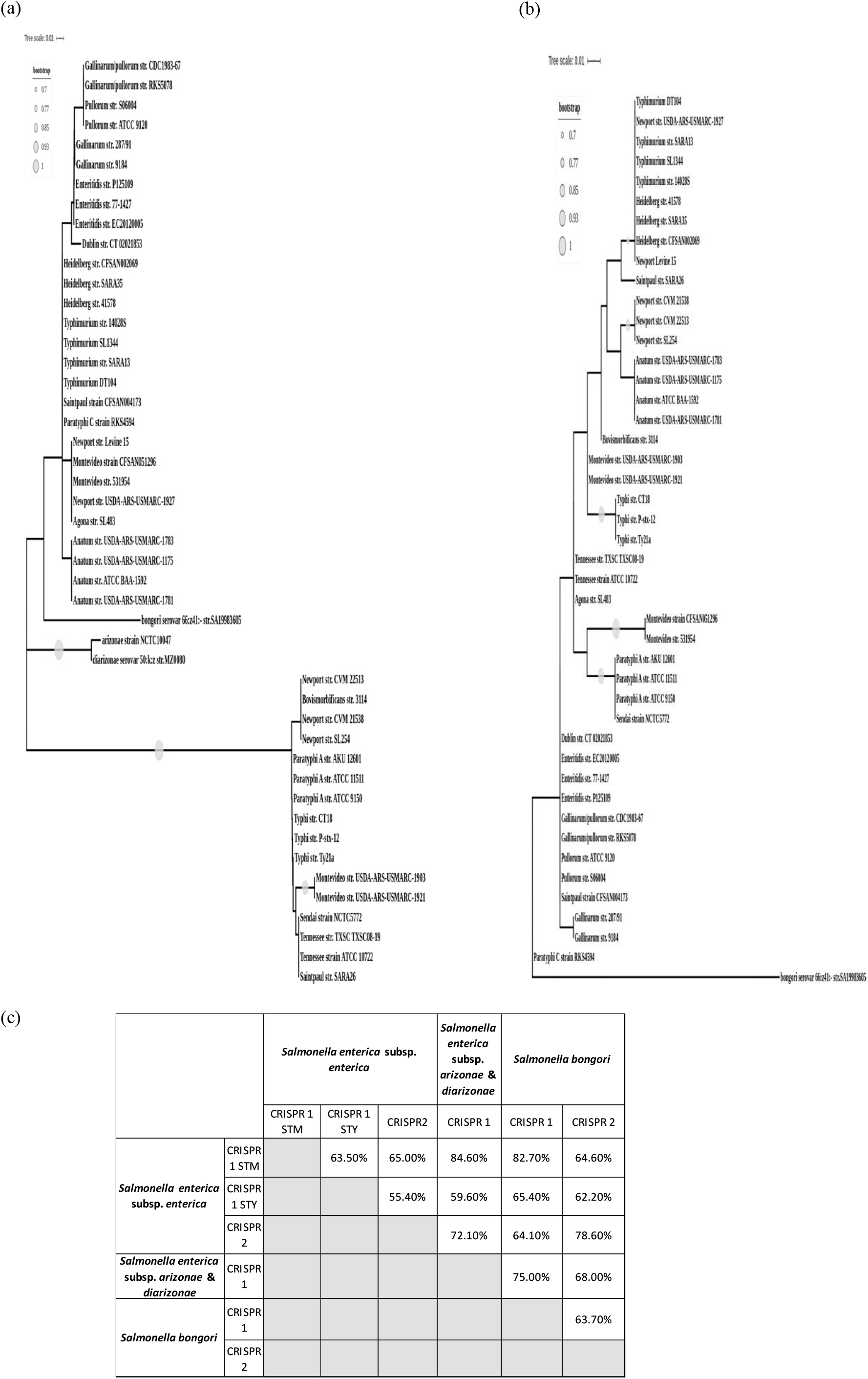
The phylogeny and conservation of CRISPR leaders. **(a & b)** Phylogeny of leaders across *Salmonella* serovars for CRISPR1 (**a**), and CRISPR2 **(b)** leaders. The CRISPR leader sequences were aligned using Muscle, and the phylogenetic trees were constructed by ML. The trees were visualised by iTOL. **(c)** The matrix depicting the inter-species and inter-subspecies conservation of the leader sequence of both the CRISPR arrays. The values represent the percent nucleotide identity with respect to the entire query cover. The reference strains are *S. enterica* serovar Typhimurium str.14028S, Typhi str. CT18, *S. enterica* subsp. *arizonae* str. NCTC10047, *S. enterica* subsp. *diarizonae* str. MZ0080 and *S. bongor*i str. SA19983605.

The serovars of *S. enterica* subsp. *enterica* have two distinct types of CRISPR1 leaders (Fig. 3 and Fig. S5), justifying their divergence in two clades. One of the leader sequence is identical to that of Newport II^11^ and is also present in all the serovars of CRISPR1-STY clade. Serovars Enteritidis and Gallinarum cluster in the CRISPR1 leader tree (Fig. 3) with 100% leader identity. Serovars Gallinarum/Pullorum and Pullorum separate from this cluster possessing an SNP (A to G transition) in the leader. On similar grounds, other serovars cluster or separate from their clustering partners. The CRISPR1 leader of *S. bongori,* and *S. enterica* subsp. *arizonae* and subsp. *diarizonae* maximally matched with that of CRISPR1-STM (Fig. 3 and Fig. S5) and hence grouped in the CRISPR1-STM clade.

The CRISPR2 leader sequence is highly conserved (with a few SNPs) among all the serovars of *S. enterica* subsp. *enterica* (Fig. 3 and Fig. S5). The variations due to SNPs explains the serovar clustering in the CRISPR2 leader tree. For instance, the leaders of serovars Paratyphi A and Typhi have 94% sequence similarity hence segregated into separate clades while the serovars Paratyphi A and Sendai clubbed together with 100% similarity.

The analyses indicate the overall topology and clustering of strains in the CRISPR1 and CRISPR2 trees is a result of differences in the leader sequence, and the spacer conservation governs the sub-clustering of the strains. The aLRT parametric assessment too coincides with the Bayesian approach (Fig. S6 & S7).

#### 2.2.4 MLST phenogram and its association with the CRISPR array

MLST is considered as a robust and widely accepted phylogenetic reconstruction of species^16^. Hence we generated a reference MLST tree for the shortlisted strains (Table S1 using concatenated allelic data of seven housekeeping genes (Fig. 4). The MLST tree topology reveals *S. bongori* to be a clear outgroup, and the most ancestral *Salmonella* lineage, consistent with the reported data^17^. All other serovars formed lineages within a serovar-specific cluster depicting to have evolved together as an individual taxon except serovar Saintpaul and Newport. In this light serovar Saintpaul turns out to be polyphyletic like serovar Newport^18^. Serovar Saintpaul str. SARA26 separated from all serovars of subspecies *enterica* while the serovar Saintpaul str. CFSAN004173 clustered with Typhimurium/Heidelberg/Newport II clade. Serovar Paratyphi A is closer to serovar Typhimurium with 98.8% similarity in the seven genes than to serovar Typhi (98.6% similarity). The CRISPR and MLST phenograms are discordant signifying independent evolution. However, serovars Montevideo-STM and Montevideo-STY possess the same housekeeping genes but differ in CRISPR arrays segregating in two groups in CRISPR1 and CRISPR2 phenograms. This analysis indicates an interlinked evolution of the CRISPR1 and the CRISPR2 arrays.

**FIGURE 4.**
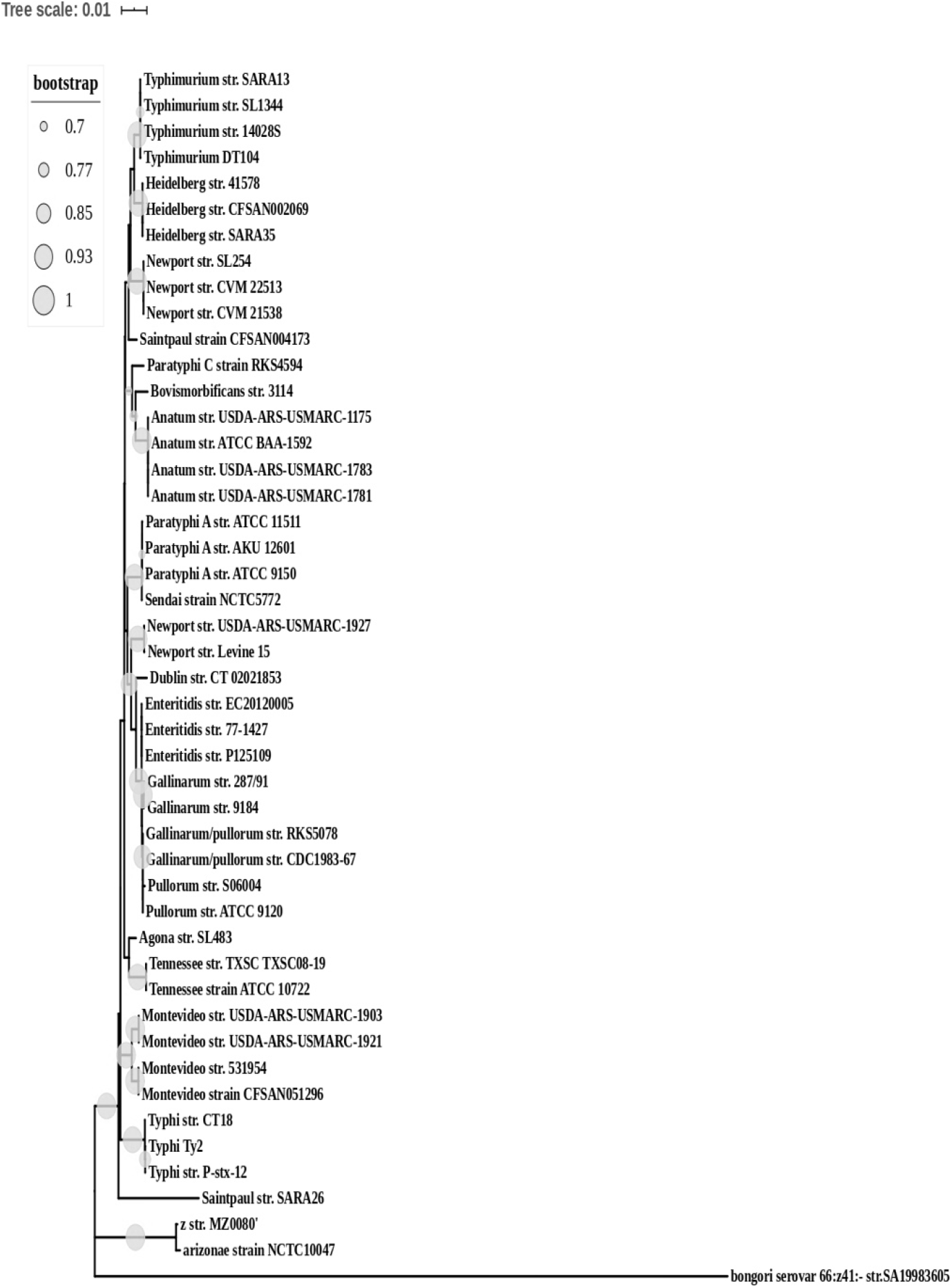
The MLST phylogeny. The phylogenetic tree was constructed using the concatenated sequence of seven housekeeping genes - *purE, hemD, aroC, dnaN, hisD, thrA,* and *sucA.* The sequences were aligned using Muscle, and the phylogenetic trees were constructed by ML. The trees were visualised by iTOL.

### 2.3 Phylogeny and classification of the *cas* operon

#### 2.3.1 Diversification of *cas* operon and its association with the CRISPR1 array

Two distinct *cas* gene arrangements were obtained for the strains comprising CRISPR1-STY and CRISPR1-STM clade. The *cas* operon of the respective categories were denoted as *cas-*STY and *cas-*STM. For *cas-*STY, the *cas3* gene is present as a complement and is singled out from the other *cas* genes by a gap of 357 nucleotides (561 for serovar Montevideo-STY) (Fig. S8). For *cas-*STM, the *cas* genes are contiguous but the *cas3* gene of serovar Montevideo-STM and *S. enterica* subsp. *arizonae* is degenerate having a premature stop codon. Moreover, we noticed structural heterogeneity within the *cas-*STM operon across CRISPR1-STM strains, with respect to its position to both the CRISPR loci and the *cas* gene composition (Fig. S8). The *cas* operon of *S. bongori, S. enterica* subsp. *enterica*, subsp. *arizonae* and subsp. *diarizonae* were termed as *cas*-STM.B, *cas*-STM.E, *cas*-STM.A, and *cas*-STM.D respectively.

#### 2.3.2 Evolutionary studies and conservation of *cas* operon in *Salmonella*

The *cas* operon’s heterogeneity was further assessed through phylogenetic analysis of the *cas3* gene and the entire *cas* operon (Fig. 5 and supplementary fig. S10). Two clades and the clustering of serovars obtained in both the phenograms is far more analogous with the CRISPR1 phenogram. To gain insights into the serovar clustering in *cas* genes, we performed a detailed comparative analysis of *cas* operon. The analysis of all *cas* genes considered in concatenation revealed the highest nucleotide similarity (99%) between subspecies *arizonae* and *diarizonae* and lowest (28.6%) between the *cas-*STM and *cas-*STY group (Fig. S9). Between *cas-*STM and *cas-*STY group, *cas1* shares the highest similarity (74.4 - 78.8% nucleotide and 82.5 - 87% amino acid match) while *cse2* shares the lowest similarity (no significant nucleotide match and 35% amino acid identity) (Fig. 5). The Cas3 nuclease differed substantially between the *cas-*STM and *cas-*STY categories (10.47 - 18.4% nucleotide and 37.4 - 45% amino acid match). However, the functionally important domains- HD domain, helicase C terminal domain, and the DEAD-box were nearly 75% similar. The *cse1* gene, was quite distinct between the *cas-*STM and *cas-*STY categories. The functionally important residues of Cse1 (*E. coli*) include Gly (157), glycine-loop residues (159-161), Lys (268), Asn (353), Glu (354) and Ala (355) required for the recognition of PAM sequences^19^ and lysine residues (289-290) for recruiting Cas3 protein^19^. Most of these residues are conserved across *cas*-STM and *cas*-STY categories (Fig. S11). This indicates that even though the Cse1 and Cas3 differs significantly between these serovars, their functionality remains conserved.

**FIGURE 5.**
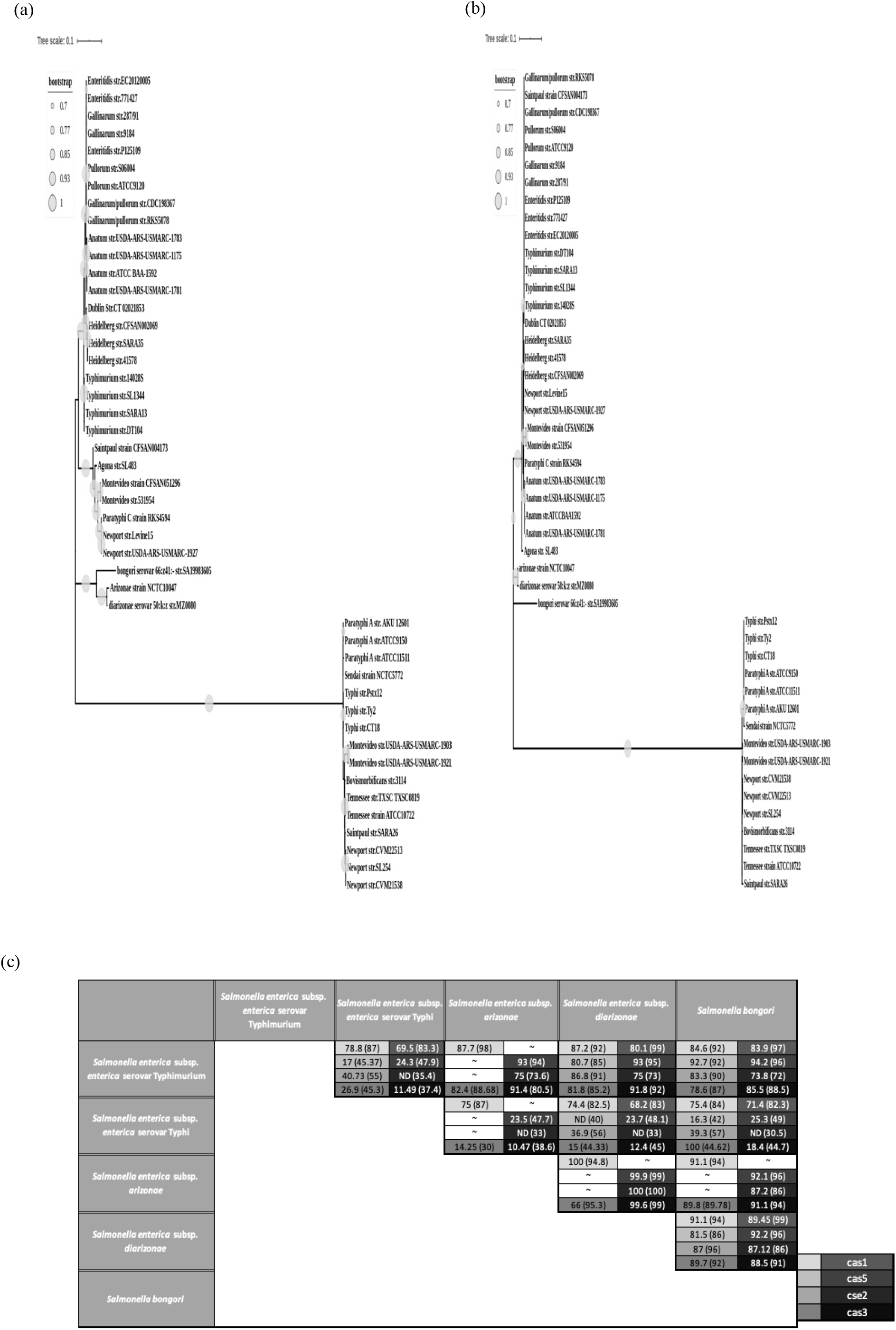
The phylogeny and conservation of *cas* genes. **(a & b)** Phylogeny of *cas* genes across *Salmonella* serovars entire *cas* operon **(a)** and the *cas3* gene **(b**). The sequences were aligned using Muscle, and the phylogenetic trees were constructed by ML. The trees were visualised by iTOL. **(c**) Conservation of all the individual *cas* gene and Cas protein sequences. The amino acid percent conservation is depicted in parenthesis. The term ‘ND’ represents no nucleotide sequence similarity based on the default parameter of the tool Nucleotide-BLAST. The reference strains are *S. enterica* serovar Typhimurium str.14028S, Typhi str. CT18, *S. enterica* subsp. *arizonae* str. NCTC10047, *S. enterica* subsp. *diarizonae* str. MZ0080 and *S. bongor*i str. SA19983605.

A noteworthy observation was made for the CRISPR1 and *cas* trees. In general, the serovars of the CRISPR1-STY/*cas*-STY clade are more evolved than those of the CRISPR1-STM/*cas*-STM clade (Fig. 2 and 5). Additionally, there is the latest evolution of the strains of Montevideo-STY compared to the other serovars. In contrast, the serovars of CRISPR1-STM/*cas*-STM clade, show limited evolution as interpreted from their scaled branching pattern. However, CRISPR2 phylogeny depicts a mixed mode of such evolution.

### 2.4 Inter-genus analysis of the CRISPR-Cas system

The evolutionary history of the CRISPR and *cas* loci across the *Enterobacteriaceae* family was studied through comparative sequence analysis and phylogenetics. Only the CRISPR1 leader of *Salmonella* showed substantial match across genera *Escherichia, Citrobacter, Shigella,* and *Klebsiella*, while the CRISPR2 leader showed match only with *Klebsiella* (75%). Thus, we constructed a CRISPR1 leader phenogram with 17 strains belonging to these genera (Table S3), and some strains of CRISPR1-STM and CRISPR1-STY clades. The phylogenetic tree diverged into two main clades (Fig. 6) similar to the CRISPR1 tree of *Salmonella* with the same signature serovars. The strains of CRISPR1-STY category grouped with *Escherichia, Shigella* and some strains of *Citrobacter* (Fig. 6 and Table S3) while the strains of CRISPR1-STM clustered with *Klebsiella,* and a strain of *Citrobacter* (Fig. 6).

**FIGURE 6.**
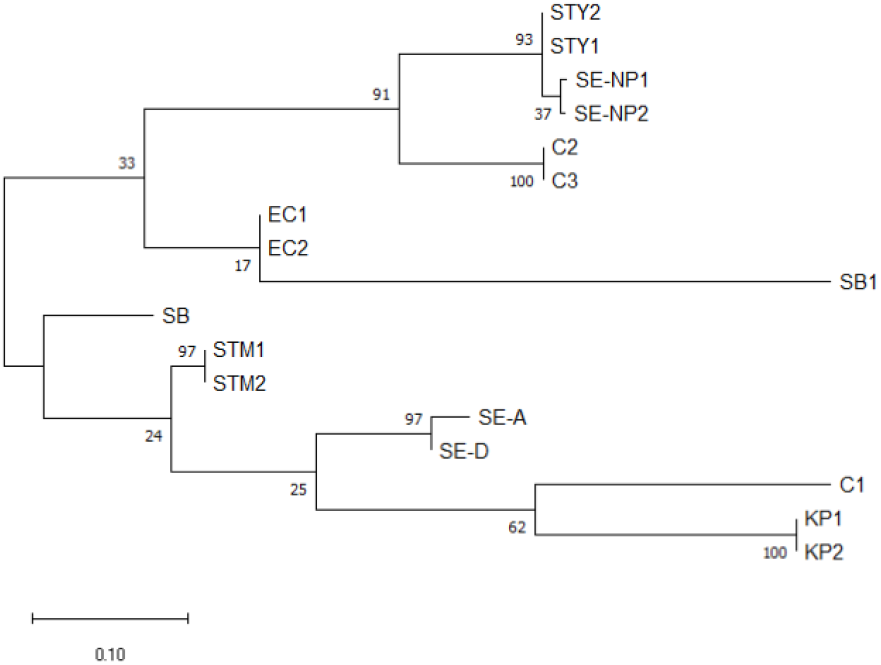
The Phylogeny of the CRISPR1 leader sequence of 17 strains of species of *Enterobacteriaceae* family. The CRISPR leader sequences were aligned using Muscle, and the phylogenetic trees were constructed by ML. The bootstrap values are indicated at each node. KP - *Klebsiella pneumoniae*, C – *Citrobacter*, SE-A – *S. enterica* subsps. *arizonae*, SE-D – *S. enterica* subsps. *diarizonae*, STM – *S. enterica* subsp. *enterica* serovar Typhimurium, STY – *S. enterica* subsp. *enterica* serovar Typhi, SE-NP *– S. enterica* subsp. *enterica* serovar Newport, SB – *S. bongori*, SB1 – *Shigella boydii* and EC – *Escherichia coli*.

The *cas-*STM operon showed ~75% similarity with that of the species*, Klebsiella pneumoniae* (str. TGH10)*, Citrobacter freundii* (sp. CFNIH3), and *Shigella boydii* (str. ATCC 49812), which is significantly higher than that with *cas*-STY (28.6%). On the contrary, the *cas-*STY operon displayed ~84% similarity with *Citrobacter freundii* (sp. CFNIH9) and *Citrobacter* (sp. 30_2). Intriguingly, the *ca*s-STY showed only a 12% match with *E. coli*.

### 2.5 CRISPR-Cas system flanked by mobile genetic elements (MGE)

To decipher the probable involvement of horizontal gene transfer (HGT), we screened the presence of the signature genes, helicase, transposase, and integrase^20,21^ in the proximity of the CRISPR-Cas region of *Salmonella* and analyzed the GC content of this region in comparison to the whole genome. We found that out of 20 serovars, 18 serovars (representative strains of each considered) showed truncated/probable transposase, 30 kb upstream of the CRISPR1 loci (Fig. 7 and Table S4). Two partial but different amino acid sequences of transposase were obtained for serovars belonging to the CRISPR1-STM and CRISPR1-STY clades. The GC content of the CRISPR arrays for most of the serovars was higher than the GC content of the whole genome except a few serovars with smaller arrays had lower GC content due to AT rich leader sequence (Fig. 7 and Table S4). A transposase gene was also present upstream of CRISPR2 array in serovars Paratyphi A and Typhi. Helicase gene was present in serovars Typhi and Typhimurium.

**FIGURE 7.**
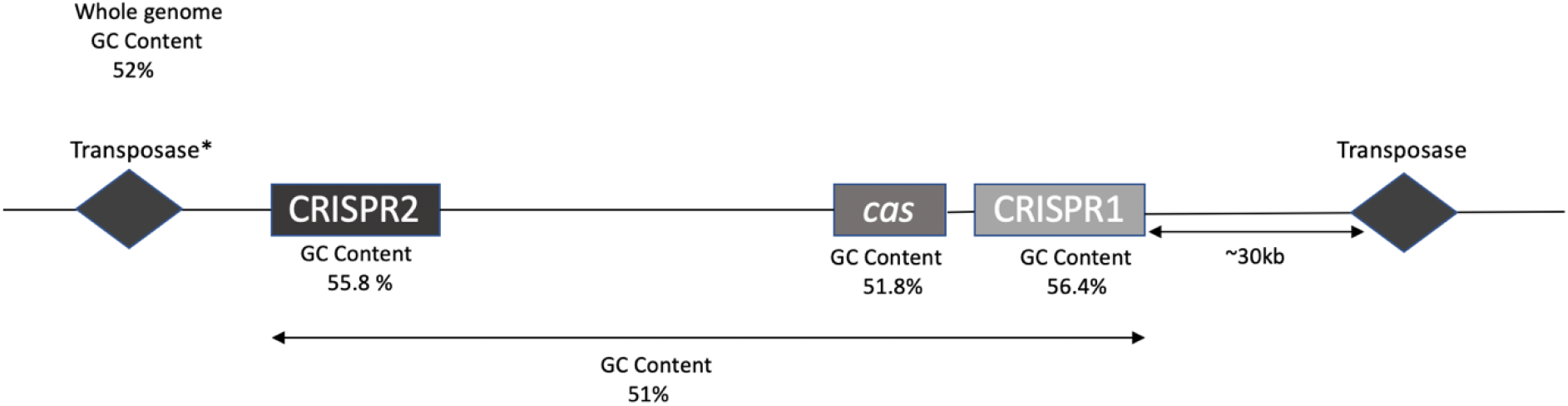
Generalised representation of the signature genes involved in horizontal gene transfer. All *Salmonella* serovars except serovars Bovismorbificans and Gallinarum/Pullorum contain the transposase gene upstream of CRISPR1 loci. * - transposase upstream of CRISPR2 is present only in serovars Typhi and Typhimurium.

## 3. DISCUSSION

The evolutionary mechanisms in bacteria are highly complex with environmental factors intricately modulating the genome architecture and functionality. This provides extra-ordinary resilience to the bacteria. When combined with HGT and recombination events the evolutionary framework of the bacteria is significantly influenced providing multiple opportunities for successful evolution as a potent pathogen. Our study depicts the evolution of *Salmonella* virulence with respect to CRISPR-Cas system that is involved in phage-immunity, pathogenicity, and genome evolution. We categorized this system into two types CRISPR1-STM/*cas*-STM and CRISPR1-STY/*cas*-STY based on the phylogenetic segregation and differences in the CRISPR1 leader and *cas* genes features of the shortlisted 47 strains. Similar segregation pattern was observed with a large set to 128 strains (Fig. S12). The CRISPR1-STM clade included strains that are host-adapted, host-specific, and have broad-host range (Fig. 2)^22,23^. The CRISPR1-STY clade comprised typhoidal serovars (Fig. 2)^24^, and serovars Montevideo, Newport II, Tennessee, Bovismorbificans and Saintpaul having broad host-range^25,26^ and association with outbreaks of human salmonellosis^27–29^.

The chronicles of battles between the bacteria and the invading MGE are registered as spacers in the CRISPR arrays. The divergence of the CRISPR-Cas system in *Salmonella* was understood through a comprehensive intra- and inter-serovar mapping of the spacer sequences. The conservation of spacers was weak across the serovars with only ~8-9.5% of unique spacers present in more than one serovar. This suggests a non-random distribution of the arrays (to be precise, spacers) foreshadowing the evolutionary kinetics of *Salmonella.* The hyper-diversity of the spacers, but their conservation within a particular serovar confers acquisition of the spacers a very primitive event with different selective pressures operating on different serovars. This could have led to a non-classic evolutionary model of the CRISPR-Cas system. This polymorphism of spacers across serotypes finds utility in serotyping^30,31^.

The CRISPR–Cas evolution is complex and the modular character hinders its forthright categorization based on the serovar host-range and geographical location. Serovars Newport II and Newport III, both infect primates, reptiles and aves but still segregate into two separate clades in the CRISPR1 tree. Serovar Typhimurium strain SARA13 and serovar Saintpaul SARA26 were isolated from the same geographic location, France (GenBank database), but segregated into CRISPR1-STM and CRISPR1-STY clades, respectively. Spacer conservation within strains of all the serovars irrespective of the geographic location suggests the roots of CRISPR origin to be ancient and the displacement has a diminutive role. Some serovars share a significant proportion of their spacers, suggesting their recent divergence in the evolutionary timeline of *Salmonella*. Example include, 66% of CRISPR1 and 100% of CRISPR2 spacers of serovar Heidelberg are identical with those of serovar Typhimurium.

The anchor spacer gives an indirect correlation of the last common ancestor (LCA) for the inter-serovar studies^11^. Most serovars of the three individual clusters in CRISPR1-STM clade of CRISPR1 tree (Fig. 2), share the anchor spacer suggesting a LCA for each. For instance, in the Enteritidis cluster serovar Enteritidis and Pullorum share anchor spacer. Serovar Gallinarum shares spacers with Enteritidis but not the anchor spacer indicating the loss of some common spacers (also by other serovars) including the anchor spacer. In the CRISPR1-STY clade, some serovars do have similar spacers, but none is the anchor spacer. For the CRISPR2 tree, even though serovars Enteritidis, Paratyphi C, Gallinarum, Pullorum, Gallinarum/Pullorum, and Typhi are part of different clusters, they have an identical anchor spacer. This suggests that the LCA for these serovars could have been communal and diverged lately due to distinct evolutionary forces. Serovar Bovismorbificans share five CRISPR1 spacers with serovar Saintpaul-STM and anchor spacer with serovar Newport II indicating divergence from Newport II and recombination with Saintpaul-STM (from CRISPR1-STM clade). Generally, the anchor spacer for a particular serovar is conserved and disparities tolerated at the leader proximal end^11^. Intriguingly, all the strains of a given serovar have a few spacers conserved, irrespective of their position in the array and the spacer variability within the serovar. For instance, in CRISPR1, spacer B in the serovar Typhimurium, and spacers A & P-R of serovar Newport III are highly conserved even though, these serovars show variations in their spacer composition (Fig. S1). One elucidation is, as implicated elsewhere^13^, the spacer composition of the system could potentially leverage the pathogenic potential possibly through endogenous gene regulation and, therefore, were retained in the system.

The *cas* genes of the strains in the *cas*-STM and *cas*-STY category are highly similar within each category but differ from the other, except for the *cas1* and *cas2,* genes required for spacer acquisition. However, the key residues of Cse1 and the functional domains of Cas3 are conserved indicating the conservation of their functionality. The strains comprising *cas*-STM and *cas*-STY are homologous to that of the CRISPR1 phenogram. This is empirical as the CRISPR1 array, and the *cas* operon lie in close proximity, suggesting an evolutionary coherence between the two. Furthermore, the strains belonging to CRISPR1-STY/*cas-*STY category showed higher substitutions per sequence site, implying the plasticity for new alterations.

The size of the spacer set for a given serovar is directly proportional to its host-range. Ubiquitous serovars like Typhimurium, Newport II, Tennessee, and Heidelberg has huge spacer set while host-specific/adapted serovars like Typhi, Sendai, Gallinarum, Dublin etc. possess a few spacers. Considering the role of spacers in regulating endogenous genes^32^, we hypothesize that versatility in spacers opens up opportunities to infect multiple hosts permitting the regulation of a variety of genes. All the spacers of the host-specific serovars Gallinarum, Pullorum, and Gallinarum/Pullorum are present in serovar Enteritidis (a broad-host-range serovar) along with some additional spacers further testifying our hypothesis. Among the host-specific/adapted serovars, the primate-specific serovars, Typhi, Paratyphi A, and Sendai, have a CRISPR1-STY leader and *cas*-STY operon. The remaining four serovars are specific to poultry or cattle containing the CRISPR1-STM/*cas*-STM type CRISPR/Cas system. With these observations, we are tempted to put forward a conjecture for *Salmonella* serovars ‘Fewer spacers in CRISPR array restrict the host-range, and the CRISPR1-STY/*cas*-STY type CRISPR/Cas system confers specificity to primate’. However, in-depth analyses and further research are warranted to understand any advantage of having a CRISPR1-STY/*cas*-STY system in infecting primates.

The incongruence in CRISPR and *cas* trees with the MLST tree implies a possible event of HGT. We tracked the presence of a truncated transposase, a hallmark of HGT, ~30 kb upstream of the CRISPR1 array with the protein sequences showing significant match within the serovars of the CRISPR1-STM (>95%) and CRISPR1-STY (88%) clades. A high GC content of the CRISPR array fortifies the occurrence of HGT event. A further support is evidenced through the histone-like nucleoid-structuring protein (H-NS) mediated regulation of *cas* operon in *S. enterica* subsp. *enterica* serovar Typhi^33^. H-NS is associated with HGT, acting as a transcriptional silencer of horizontally acquired genes by binding to the AT rich DNA and blocking RNA polymerase^3^. One may possibly argue the regulation of CRISPR array by H-NS through its AT-rich leader. With this background, we anticipate that H-NS could have originally silenced the CRISPR-Cas system and later evolved to regulate the functioning of *cas* operon and probably the CRISPR arrays. It is quite likely that all the strains belonging to CRISPR1-STY clade conform to this mechanism. While validation of such mechanism in the strains of CRISPR1-STM clade needs accreditation.

The CRISPR and *cas* attributes of *S. bongori*, *S. enterica* subsp. *arizonae* and subsp. *diarizonae,* justify their placement in ‘CRISPR1-STM’ and ‘*cas*-STM’ clades of CRISPR1 and the *cas* phenograms, thereby reflecting a higher similarity with CRISPR1-STM/*cas*-STM (~85%) than with CRISPR1-STY/*cas*-STY (20-30%). Surprisingly, the CRISPR1-STM/*cas*-STM and CRISPR1-STY/*cas*-STY sequences showed better relatedness with other genera of *Enterobacteriaceae* family namely, *Escherichia*, *Klebsiella, Shigella,* and *Citrobacter* than with each other. A significant proportion of the *Escherichia, Shigella,* and *Klebsiella* strains have the CRISPR/Cas system that matched with CRISPR1-STM/*cas*-STM (unpublished data). Nevertheless, few strains of the *Enterobacteriaceae* family (*Klebsiella* & *Citrobacter*) contain both the type of CRISPR1 array and *cas* operon. Moreover, in *Citrobacter freundii* complex sp. CFNIH3, a truncated transposase was found 30 kb upstream of the CRISPR1 loci with a 40% similarity in the connecting-region of *S. enterica* subsp. *enterica* serovar Typhimurium indicating occurrence of HGT event. The split of *Salmonella* serovars into two separate clades and clubbing of serovar of CRISPR1-STM with *Shigella* and *E. coli* was also observed in the Cas2 protein phylogram reported by Touchan *et al.*^12^ thus, validating our results.

With the comprehensive analysis of all the results, we put forward the following hypotheses for evolution of CRISPR-Cas system. Given that a significant proportion of *Escherichia, Shigella,* and *Klebsiella* strains harbor CRISPR1-STM/*cas*-STM type leader and operon, we propose that the LCA of the *Enterobacteriaceae* family could have had the CRISPR1-STM/*cas*-STM type loci. Further, after the divergence from *Escherichia*, *Salmonella* could have acquired its CRISPR2 array, as there exists no similarity in their leader sequences, while leaders of *S. enterica* and *S. bongori* are 78% identical. *S. enterica* subsp. *arizonae* and subsp. *diarizonae* do not have a CRISPR2 array, which could have been possibly lost in due course of evolution. Many strains of subsp. *arizonae* do not contain the CRISPR1 array suggesting its loss as well. We also observed the conservation of CRISPR2 leader throughout *S. enterica subsp. enterica*, the existence of similar spacers of CRISPR2 array across the strains, and no bifurcation in the CRISPR2 phenograms. With this background, we propose the following. Apparently one, few or all the serovars belonging to the CRISPR1-STY/*cas*-STY clade could have acquired CRISPR1-STY leader and *cas*-STY operon from an unknown source, possibly by HGT event in the gut of primates, subsequently transmitting amongst other *Salmonella* strains and other genera whereas the CRISPR2 array remained unaffected. The inheritance of the CRISPR1-STY/*cas*-STY system perhaps renders competitive advantage to the strains possessing it, in terms of its pathogenicity and enhanced survival in hostile conditions. Further investigation of CRISPR-Cas evolution across *Enterobacteriaceae* family is warranted to strengthen our hypothesis.

The results of the study hold prospect in providing insights into the evolution of *Salmonella* (other enteropathogens) as a pathogen with diverse host-specificity, linking various regulatory networks with the CRISPR-Cas system.

## 4. MATERIALS AND METHODS

### 4.1 Sequence Data Collection

The CRISPR and *cas* loci of 131 *Salmonella* strains were obtained in correct orientation after retrieving the data from GenBank and CRISPR-Cas++ database^34^. For details, refer to supplementary material. For MLST, sequences of seven housekeeping genes namely*, purE, hemD, aroC, dnaN, hisD, thrA* and *sucA* were retrieved from BIGSdb software^35^, and the unannotated ones were extracted from the genome’s annotation files using customized written bash script. The composite sequence tags were allocated for the allelic profiles of these seven genes.

The CRISPR leader sequences of 17 strains comprising of genus *Salmonella, Escherichia, Citrobacter, Shigella, and Klebsiella* were extracted using the CRISPR-Cas++ database/CRISPRCasFinder and matched with the *Salmonella’s* CRISPR leader sequences.

### 4.2 Analysis of the CRISPR-Cas components

To create spacer maps of the CRISPR arrays, the spacers were aligned to maximize their homology across the *Salmonella* strains. The intra- and inter-serovar spacer conservation were estimated using python scripts. The orientation of the individual *cas* genes was traced and the sequence similarity calculated using a custom python script. The amino acid sequences of Cse1 and the essential domains of Cas3 protein (HD domain, helicase C terminal domain, and the DEAD-box) of *Salmonella* were extracted from the Uniprot database and aligned with the reported sequences of *E. coli* using the tool Clustal Omega.

The sequence logo for the CRISPR leader and DR sequences^34^ were generated using the tool WEBLOGO ver 2.8.2^43^. The MGEs elements were manually checked 50 kb upstream and downstream of each CRISPR loci using the annotated GenBank files. Further, the GC content of the CRISPR-Cas components, and the whole genome was computed using python script.

### 4.3 Phylogenetic analysis

Multiple sequence alignment were performed on the aforesaid sequences by MUSCLE version 3.6 with default parameters^36^ integrated into Molecular Evolutionary Genetics Analysis version 10 (MEGA X)^37^. All positions with alignment gaps and missing data were excluded (complete deletion option). The resulting alignments of respective groups of sequences were used to construct each phylogenetic tree using the Maximum Likelihood (ML) method^38^ guided by the most suitable evolutionary model proposed by Bayesian approach^39^. The trees were given confidence with a bootstrap value of 1000 iterations. The substitution models and the parameters used for the reconstructed trees were Tamura-Nei model with Gamma distribution for MLST; Tamura 3-parameter model for CRISPR1 leader and CRISPR2 Leader; Kimura 2-parameter model for CRISPR1 leader with array and CRISPR2 leader with array; and Kimura 2-parameter model along with gamma distribution for concatenated *cas* genes and *cas3* gene. The Newick format of the trees was used for further visualization and analyses through MEGA X and iTOL^40^. All trees were drawn to scale, and the branch lengths were calculated as the number of substitutions per site.

For topology validation, the phylogenetic trees for *Salmonella* were also reconstructed using the program PHYML version 3.1^41^ with statistical tests for branch support. The statistical parametric analysis of Shimodaira-Hasegawa re-estimation of log-likelihood was adopted to get the consensus maximum likelihood tree. The general time reversible substitution models were kept uniform for all the trees generated. Curation of the multiply aligned sequences was done through GBlocks, having, a stringent selection of many contiguous non-conserved positions being disallowed^42^.

## Acknowledgments

This work was supported by the Department of Science and Technology, Science and Engineering Research Board (Grant No. ECR_2017_002053) to SAM. The authors acknowledge the support of the Department of Biological Sciences, Sunway University, Selangor, Malaysia for providing the computational facilities.

## Author contributions

S.K.K initiated the research and completed the main work under the S.A.M and C.L. guidance. B.A. carried out the computational assessments. S.A.M finalized the paper. All authors reviewed the manuscript.

## Competing interests

The authors declare they have no competing interests.

## Notes

### Competing Interest Statement

The authors have declared no competing interest.

