## Supplemental file for "The Phylogenomics of CRISPR-Cas System in *Salmonella*: an evolutionary race with housekeeping genes"

### Supplementary material for “The Phylogenomics of CRISPR-Cas System in *Salmonella*: an evolutionary race over housekeeping genes” by Kushwaha et al.

**Supplementary materials include:**

**Table S1-S4**

**Supplementary figure legends**

**Figures S1-S12**

**Supplementary Methodology**

**Supplementary Table S1. Name and accession number for whole genome sequence for the strains analysed in the study.**

| STRAIN NAME | ACCESSION NUMBER | STRAIN NAME | ACCESSION NUMBER |
|---|---|---|---|
| *Salmonella enterica* subsp. *enterica* | | | |
| Serovar Anatum | | Serovar Paratyphi A | |
| **USDA-ARS-USMARC-1175** | **CP007483.2** | **AKU_12601** | **FM200053.1** |
| USDA-ARS-USMARC-1676 | CP014620.1 | **ATCC 11511** | **CP019185.1** |
| USDA-ARS-USMARC-1677 | CP014663.2 | **ATCC 9150** | **CP000026.1** |
| USDA-ARS-USMARC-1727 | CP014621.2 | Serovar Gallinarum | |
| USDA-ARS-USMARC-1728 | CP014664.1 | **287/91** | **AM933173.1** |
| USDA-ARS-USMARC-1735 | CP007584.2 | **9184** | **CP019035.1** |
| USDA-ARS-USMARC-1736 | CP014657.1 | Serovar Pullorum | |
| USDA-ARS-USMARC-1765 | CP014659.2 | **ATCC 9120** | **CP012347.1** |
| USDA-ARS-USMARC-1766 | CP014665.1 | **S06004** | **CP006575.1** |
| **USDA-ARS-USMARC-1781** | **CP014666.2** | Serovar Gallinarum/Pullorum | |
| **USDA-ARS-USMARC-1783** | **CP014661.1** | **CDC1983-67** | **CP003786.1** |
| **ATCC BAA-1592** | **CP007531.1** | **RKS5078** | **CP003047.1** |
| CDC 06-0532 | CP007211.2 | Serovar Typhimurium | |
| Serovar Enteritidis | | USDA-ARS-USMARC-1808 | CP014969.1 |
| **77-1427** | **CP007598.1** | USDA-ARS-USMARC-1810 | CP014982.2 |
| CDC_2010K_0968 | CP007528.1 | USDA-ARS-USMARC-1880 | CP014981.1 |
| EC20090641 | CP007249.2 | USDA-ARS-USMARC-1896 | CP014977.1 |
| EC20090698 | CP007248.1 | USDA-ARS-USMARC-1898 | CP014971.2 |
| EC20110221 | CP007247.1 | USDA-ARS-USMARC-1899 | CP007235.2 |
| EC20110354 | CP007175.1 | **str. 14028S** | **CP001363.1** |
| EC20110355 | CP007250.1 | str. 798 | CP003386.1 |
| EC20110356 | CP007262.1 | CDC 2009K-1640 | CP014975.1 |
| EC20110357 | CP007261.1 | CDC 2009K-2059 | CP014983.1 |
| EC20110358 | CP007260.1 | CDC 2010K-1587 | CP014965.1 |
| EC20110360 | CP007258.1 | CDC 2011K-0870 | CP007523.1 |
| EC20110361 | CP007263.1 | CDC 2011K-1702 | CP014967.1 |
| EC20111095 | CP007254.1 | D23580 | LS997973.1 |
| EC20111174 | CP007253.1 | **DT104** | **HF937208.1** |
| EC20111175 | CP007252.1 | DT2 | HG326213.1 |
| EC20120002 | CP007329.2 | L-3553 | AP014565.1 |
| **EC20120005** | **CP007267.2** | **SARA13** | **CP017728.1** |
| EC20120008 | CP007245.1 | **SL1344** | **FQ312003.1** |
| EC20121175 | CP007269.2 | ST4/74 | CP002487.1 |
| **P125109** | **AM933172.1** | T000240 | AP011957.1 |
| Serovar Saintpaul | | UK-1 | CP002614.1 |
| **CFSAN004173** | **CP019204.1** | CFSAN001921 | CP006048.1 |
| CFSAN004174 | CP019206.1 | Serovar Sendai | |
| CFSAN004175 | CP019172.1 | **NCTC5772** | **NZ_UGVR01000002.1** |
| SGB23 | CP023166.1 | Serovar Tennessee | |
| **SARA26** | **CP017727.1** | **ATCC 10722** | **CP025218.1** |
| Serovar Heidelberg | | CFSAN001387 | CP014994.1 |
| CFSAN002064 | CP005995.1 | CFSAN070643 | CP024168.1 |
| **41578** | **CP004086.1** | CFSAN076210 | CP033345.1 |
| B182 | CP003416.1 | PIR00537 | CP025217.1 |
| **CFSAN002069** | **CP005390.2** | CFSAN070645 | CP024164.1 |
| **SARA35** | **CP019176.1** | **TXSC_TXSC08-19** | **CP007505.1** |
| SL476 | CP001120.1 | Serovar Dublin | |
| Serovar Typhi | | **CT_02021853** | **CP001144.1** |
| **P-stx-12** | **CP003278.1** | Serovar Bovismorbificans | |
| **Ty2** | **AE014613.1** | **3114** | **HF969015.2** |
| **CT18** | **AL513382.1** | Serovar Paratphi C | |
| 1016889 | LT904893.1 | **RKS4594** | **CP000857.1** |
| 1036491 | LT904885.1 | Serovar Montevideo | |
| 129-0238-M | LT904888.1 | **CFSAN051296** | **CP029336.1** |
| ERL034151 | LT904867.1 | **531954** | **CP029035.1** |
| Serovar Newport II | | 507440-20 | CP007530.1 |
| **CVM 21538** | **CP010282.1** | CDC 07-0954 | CP017974.1 |
| CVM 21550 | CP010283.1 | CDC 08-1942 | CP017975.1 |
| CVM 22425 | CP010279.1 | CDC 2010K-0257 | CP020912.1 |
| CVM 22462 | CP010280.1 | CDC 2011K-1674 | CP017976.1 |
| **CVM 22513** | **CP010281.1** | CDC 2012K-1544 | CP017977.1 |
| CVM N1543 | CP010284.1 | CDC 2013K-0218 | CP017978.1 |
| CVM N18486 | CP009561.1 | FCC0123 | CP040379.1 |
| **SL254** | **CP001113.1** | CDC 2009K-0792 | CP020752.1 |
| USMARC-S3124.1 | CP006631.1 | USDA-ARS-USMARC-1900 | CP017970.1 |
| WA_14882 | CP016357.1 | USDA-ARS-USMARC-1901 | CP017971.1 |
| Serovar Newport III | | USDA-ARS-USMARC-1904 | CP017972.1 |
| CFSAN001660 | CP016010.1 | USDA-ARS-USMARC-1912 | CP017973.1 |
| CDC 2009K-1331 | CP025248.1 | **USDA-ARS-USMARC-1903** | **CP007222.1** |
| CDC 2012K-0938 | CP025246.1 | **USDA-ARS-USMARC-1921** | **CP007540.2** |
| Levine 1 | CP015923.1 | *Salmonella enterica subsps. arizonae* | |
| **USDA-ARS-USMARC-1927** | **CP007216.2** | **NCTC10047** | **LR134156.1** |
| **Levine 15** | **CP015924.1** | *Salmonella enterica subsps. diarizonae* | |
| Serovar Agona | | **50 MZ0080** | **CP022142.1** |
| **SL483** | **CP001138.1** | *Salmonella bongori* | |
| | | **66 SA19983605** | **CP022120.1** |

The strains represented in bold were considered for further analysis. The strains highlighted in grey for serovars Montevideo and Saintpaul represent the ones having distinct spacer sets.

**Supplementary Table S2. The statistics of the spacer index for the serovars under consideration.**

| | | CRISPR1 | | | CRISPR2 | | |
|---|---|---|---|---|---|---|---|
| | | Minimum | Maximum | Average | Minimum | Maximum | Average |
| *Salmonella enterica* subps. *enterica* | Anatum | 2 | 8 | 4.77 | 8 | 26 | 20.25 |
| | Typhi | 6 | 7 | 6.14 | 1 | 1 | 1 |
| | Paratyphi A | 5 | 7 | 6.33 | 3 | 3 | 3 |
| | Gallinarum | 2 | 2 | 2 | 10 | 10 | 10 |
| | Pullorum | 2 | 2 | 2 | 2 | 6 | 4 |
| | Gallinarum/Pullorum | 2 | 2 | 2 | 6 | 6 | 6 |
| | Sendai | 4 | 4 | 4 | 1 | 1 | 1 |
| | Heidelberg | 25 | 25 | 25 | 16 | 18 | 17.67 |
| | Newport II | 17 | 26 | 24.4 | 12 | 19 | 18.3 |
| | Newport III | 4 | 18 | 11.33 | 10 | 20 | 16.83 |
| | Enteritidis | 8 | 9 | 8.95 | 8 | 12 | 11.35 |
| | Typhimurium | 8 | 28 | 17.95 | 15 | 35 | 25.39 |
| | Tennessee | 41 | 63 | 52.43 | 21 | 23 | 22.14 |
| | Montevideo | 4 | 36 | 23.59 | 16 | 25 | 19.94 |
| | Saintpaul | 11 | 41 | 19.4 | 20 | 22 | 20.8 |
| | Agona | 18 | 18 | 18 | 8 | 8 | 8 |
| | Paratyphi C | 10 | 10 | 10 | 9 | 9 | 9 |
| | Dublin | 2 | 2 | 2 | 5 | 5 | 5 |
| | Bovismorbificans | 24 | 24 | 24 | 15 | 15 | 15 |
| *Salmonella enterica subsp. arizonae* | | 23 | 23 | 23 | - | - | - |
| *Salmonella enterica* subsp. *diarizonae* | | 29 | 29 | 29 | - | - | - |
| *Salmonella bongori* | | 20 | 20 | 20 | 17 | 17 | 17 |

**Supplementary Table S3. Name and accession number for whole genome sequence for the strains analysed in the study.**

| SPECIES | ACCESSION NUMBER | ABBREVATION |
|---|---|---|
| *Salmonella bongori* str. 66 SA19983605 | CP022120.1 | SB |
| *Salmonella enterica* subsps. *arizonae* str. NCTC10047 | LR134156.1 | SE-A |
| *Salmonella enterica* subsps. *diarizonae* 50 MZ0080 | CP022142.1 | SE-D |
| *Salmonella enterica* subsp. *enterica* serovar Typhimurium str. 14028s | CP001363.1 | STM1 |
| *Salmonella enterica* subsp. *enterica* serovar Typhimurium str. SARA13 | CP017728.1 | STM2 |
| *Salmonella enterica* subsp. *enterica* serovar Typhi str. Ty2 | AE014613.1 | STY1 |
| *Salmonella enterica* subsp. *enterica* serovar Typhi str. CT18 | AL513382.1 | STY2 |
| *Salmonella enterica* subsp. *enterica* serovar Newport str. CVM21538 | CP010282.1 | SE-NP1 |
| *Salmonella enterica* subsp. *enterica* serovar Newport str. CVM22513 | CP010281.1 | SE-NP2 |
| *Klebsiella pneumoniae* subsp. *pneumoniae* strain TGH10 | CP012744.1 | KP1 |
| *Klebsiella pneumoniae* strain INF235-sc-2280127 | CP031817.1 | KP2 |
| *Shigella boydii* strain ATCC 49812 | CP026836.1 | SB1 |
| *Citrobacter freundii* complex sp. CFNIH3 | CP026235.1 | C1 |
| *Citrobacter freundii* complex sp. CFNIH9 | CP026238.1 | C2 |
| *Citrobacter* sp. 30_2 | CP022311.1 | C3 |
| *Escherichia coli* strain 52148 | CP050382.1 | EC1 |
| *Escherichia coli* strain SCU-118 | CP051716.1 | EC2 |

The CRISPR1 leader sequences of these strains were used for the phylogenetic analysis. The abbreviations are the key to the figure.

**Supplementary Table S4. MGE candidates flanking the CRISPR-Cas system.**

| | Genome location (Loci start and Loci end) | | | MGE | Percentage GC Content | | | | |
|---|---|---|---|---|---|---|---|---|---|
| | CRISPR2 | *cas* | CRISPR1 | Transposase/ Helicase | CRISPR2 loci | *cas* operon | CRISPR1 loci | CRISPR-Cas | Whole genome |
| Paratyphi A str. AKU_12601 | 2902111-2902322 | 2885645-2894598 | 2885105-2885560 | 2856664-2857091 and 3007579-3008037 | 51.3* | 49.9 | 53.2* | 49 | 52.18 |
| Newport II str. SL254 | 3073142-3074328 | 3056558-3064452 | 3054859-3056473 | 3024723-3025160 | 54.2 | 50.4 | 55.6 | 51 | 52.22 |
| Newport III str. USDA-ARS-USMARC-1927 | 975001-975639 | 983339-991789 | 991886-993012 | 1020178-1020414 | 51.3 | 52 | 57.2 | 51 | 52.18 |
| Heidelberg str. SL476 | 3069137-3070263 | 3052976-3061435 | 3051217-3052879 | 3022734-3022874 | 56.3 | 53.1 | 56.2 | 53 | 52.07 |
| Enteritidis str. EC20121175 | 2967508-2968269 | 2951453-2959906 | 2950779-2951356 | 2923727-2923816 | 55.8 | 52.8 | 55.7 | 51 | 52.17 |
| Gallinarum str. 9184 | 1224776-1225415 | 1233429-1241471 | 1241469-1241716 | 1271625-1271848 | 56.8 | 53.3 | 47.6* | 51 | 52.2 |
| Pullorum str. ATCC 9120 | 3871330-3871722 | 3855356-3863734 | 3855036-3855283 | 3827981-3828070 | 53.6* | 52.9 | 48.4* | 51 | 52.19 |
| Gallinarum/Pullorum str. CDC1983-67 | 2947545-2947937 | 2931570-2939948 | 2931250-2931497 | | 53.6* | 52.9 | 48.4* | 51 | 52.23 |
| Montevideo-STY str. USDA-ARS-USMARC-1900 | 1003049-1004420 | 1011949-1021097 | 1021182-1023406 | 1050505-1050672 | 57.7 | 50.4 | 57 | 52 | 52.35 |
| Montevideo-STM str. CDC 2010K-0257 | 992948-994014 | 1010602-1010052 | 1010149-1011641 | 1038743-1038910 | 56.6 | 52.4 | 55.9 | 51 | 52.21 |
| Bovismorbificans str. 3114 | 2976895-2977839 | 2960410-2969354 | 2958839-2960331 | | 54 | 50 | 55.6 | 50 | 52.16 |
| Anatum str. CDC 06-0532 | 970192-971440 | 979153-987612 | 987709-988225 | 1018009-1018429 | 56.2 | 52.8 | 55.5 | 51 | 52.18 |
| Tennessee str. ATCC 10722 | 963889-965320 | 972914-981858 | 981943-985815 | 1015542-1015960 | 59.3 | 50.4 | 57.1 | 52 | 52.23 |
| Saintpaul-STM str. CFSAN004173 | 946615-947863 | 955563-964025 | 964122-965066 | 993235-993458 | 57.2 | 52 | 57.4 | 51 | 52.21 |
| Saintpaul-STY str. SARA26 | 944744-946114 | 953779-962723 | 962808-965337 | 993600-993740 | 58.2 | 50.3 | 56.5 | 51 | 52.05 |
| Dublin str. CT_02021853 | 3137409-3137742 | 3121348-3129807 | 3121100-3121350 | 3090967-3091107 | 51.6* | 53 | 44.6* | 51 | 52.18 |
| Agona str. SL483 | 3005517-3006033 | 2989328-2997778 | 2988105-2989231 | 2956649-2956789 and 2975984-2977192 | 54.2 | 52 | 56.3 | 51 | 52.08 |
| Typhimurium str. CFSAN001921 | 473159-474652 | 482228-490687 | 490784-492564 | 522104-522291 and 423309-425144 | 55.9 | 50.4 | 56.9 | 51 | 52.17 |
| Typhi str. CT18 | 2943208-2943716 | 2926652-2935104 | 2926182-2926567 | 2898592-2899034 and 3013615-3014073 | 39.8* | 50.4 | 57.3 | 50 | 52.05 |
| Subsp. *arizonae* | | 962736-971156 | 971253-972682 | | | 52.9 | 57.4 | 52 | 51.38 |
| Subsp. *diarizonae* | | 1126557-1134989 | 1135086-1136883 | | | 52.9 | 56.7 | 53 | 51.54 |
| *S. bongori* | 922139-923204 | 930621-939089 | 939186-940434 | | 53.4 | 52.4 | 54.5 | 50 | 51.33 |

*The lower GC content of CRISPR arrays due to the AT-rich leader sequences are represented by asterisks

#### SUPPLEMENTARY FIGURE LEGENDS

**Fig. S1 Spacer conservation of various serovars of *Salmonella*.** Spacer conservation in **A.** CRISPR1 array and **B.** CRISPR2 array across *Salmonella* serovars. **C&D.** Inter-serovar spacer conservation in both, **C** the CRISPR1 and **D** CRISPR2 arrays. The colour code for a particular column represents spacer sequences with greater than 92%. The number denotes the position of the spacer from the leader sequence. The direct repeats have been eliminated for simplicity. The asterisk (*) represents the host-specific serovars. The duplication and triplication is depicted as a pattern in the coloured box.

**Fig. S2. The sequence logo of the consensus DR sequence and last direct repeat.** The DR sequence of all the strains belonging to the CRISPR1-STM, CRISPR1-STY and CRISPR2 were considered and the Web-logo of these were generated. The last direct repeat (distal to the leader sequence) was observed to contain SNPs. **A.** CRISPR1-STM consensus DR**, B.** CRISPR1-STM last DR**, C.** CRISPR1-STY consensus DR**, D.** CRISPR1-STY last DR**, E.** CRISPR2 consensus DR, and **F.** CRISPR2 last DR

**Fig. S3. The Phylogenetic verification of CRISPR1 leader and array**. The statistical validation of the CRISPR1 leader and array tree. The values indicate the aLRT scores.

**Fig. S4. The Phylogenetic verification of CRISPR2 leader and array**. The statistical validation of the CRISPR2 leader and array tree. The values indicate the aLRT scores.

**Fig. S5. The sequence logos of the leader sequence for each CRISPR array.** The leader sequence of all the strains were aligned and a web-logo was generated. The CRISPR1 leader sequence of *Salmonella enterica* subsp. *enterica* is of two types and is represented as CRISPR1-STM **(A-C)** and CRISPR1-STY **(D)**. The CRISPR2 **(E-F)** leader sequence is present and highly conserved among all the serovars except *Salmonella enterica* subsp. *arizonae* and subps. *diarizonae* . All the three sequences are also highly conserved. **A.** CRISPR1-STM *S. enterica* subsp. *enterica,* **B**. CRISPR1 *S. enterica* subsp. *arizonae* and subsp. *diarizonae enterica* **C.** CRISPR1 *Salmonella bongori* **D.** CRISPR1-STY *S. enterica* **E.** CRISPR2 *S. enterica* subsp. *enterica.* **F**. CRISPR2 *Salmonella bongori*

**Fig. S6. The Phylogenetic verification of CRISPR1 leader**. The statistical validation of the CRISPR1 leader tree. The values indicate the aLRT scores.

**Fig. S7. The Phylogenetic verification of CRISPR2**. The statistical validation of the CRISPR2 tree. The values indicate the aLRT scores.

**Fig. S8. Orientation of the CRISPR array and the *cas* operon in *Salmonella.*** Five types of arrangements were evident in *Salmonella*. The *cas*-STY arrangement present in strains belonging to the CRISPR1-STY clade. The *cas-STM* type operon were subdivided into four types- *S. enterica* subsp. *enterica* (*cas*-STM, belong to CRISPR1-STM clade), *S. bongori (cas*-STM.B), *S. enterica* subsp. *enterica*, subsp. *arizonae* (*cas*-STM.A) and subsp. *diarizonae* (*cas*-STM.D) All the strains of serovar Montevideo-M and *S. enterica* subps. *diarizonae* str. MZ0080 (used in our study) contain a non-sense mutation in *cas3* and is represented by an asterisk (*). *Salmonella bongori* str. SA19983605 (used in our study) does not contain the *cas7* gene and is depicted by a hash (#) and all strains of *S. enterica* subsp. *arizonae* contain a stop codon in the *cas3* operon represented by a thunderbolt.

**Fig. S9. The percentage conservation of the entire *cas* operon.** The values in the matrix represent the percentage nucleotide match between the categories. The reference strains are *S. enterica* serovar Typhimurium str.14028S, Typhi str. CT18, *S. enterica* subsp. *arizonae* str. NCTC10047, *S. enterica* subsp. *diarizonae* str. MZ0080 and *S. bongor*i str. SA19983605.

**Fig. S10. The Phylogenetic verification of *cas3* gene sequence.** The statistical validation of the *cas3* tree. The values indicate the aLRT scores.

**Fig. S11. The multiple sequence alignment of Cse1 protein by Clustal Omega.** The bold red color represents fully conserved residues (*) and strongly similar groups with a score >0.5 in Gonnet PAM 250 matrix (:) and the red color represent groups of weekly similar properties with a score <0.5 (.) for the residues essential for PAM recognition. The residues essential for Cas3 protein recruitment is indicated by the red box.

**Fig. S12. The Phylogeny of CRISPR1 leader**. The CRISPR1 leader sequences of 128 strains were aligned using Muscle and the phylogenetic trees was constructed by Maximum Likelihood.

### Figure S1

A

| | | | | | | | | | | | | | | | | | | |
|---|---|---|---|---|---|---|---|---|---|---|---|---|---|---|---|---|---|---|
| SL483 | 1 | 2 | 3 | 4 | 5 | 6 | 7 | 8 | 9 | 10 | 11 | 12 | 13 | 14 | 15 | 16 | 17 | 18 |
| AGONA | A | B | C | D | E | F | G | H | I | J | K | L | M | N | O | P | Q | R |

| | | | | |
|---|---|---|---|---|
| NCTC5772 | 1 | 2 | 3 | 4 |
| * SENDAI | A | B | C | D |

| | | | | | | | | | | |
|---|---|---|---|---|---|---|---|---|---|---|
| RKS4594 | 1 | 2 | 3 | 4 | 5 | 6 | 7 | 8 | 9 | 10 |
| PARATYPHI C | A | B | C | D | E | F | G | H | I | J |

| | | | | | | | | | |
|---|---|---|---|---|---|---|---|---|---|
| 77-1427 | 1 | 2 | 3 | 4 | 5 | 6 | 7 | 8 | 9 |
| CDC_2010K_0968 | 1 | 2 | 3 | 4 | 5 | 6 | 7 | 8 | 9 |
| EC20090641 | 1 | 2 | 3 | 4 | 5 | 6 | 7 | 8 | 9 |
| EC20090698 | 1 | 2 | 3 | 4 | 5 | 6 | 7 | 8 | 9 |
| EC20110221 | 1 | 2 | 3 | 4 | 5 | 6 | 7 | 8 | 9 |
| EC20110354 | 1 | 2 | 3 | 4 | 5 | 6 | 7 | 8 | 9 |
| EC20110355 | 1 | 2 | 3 | 4 | 5 | 6 | 7 | 8 | 9 |
| EC20110356 | 1 | 2 | 3 | 4 | 5 | 6 | 7 | 8 | 9 |
| EC20110357 | 1 | 2 | 3 | 4 | 5 | 6 | 7 | 8 | 9 |
| EC20110358 | 1 | 2 | 3 | 4 | 5 | 6 | 7 | 8 | 9 |
| EC20110360 | 1 | 2 | 3 | 4 | 5 | 6 | 7 | 8 | 9 |
| EC20110361 | 1 | 2 | 3 | 4 | 5 | 6 | 7 | 8 | 9 |
| EC20111095 | 1 | 2 | 3 | 4 | 5 | 6 | 7 | 8 | 9 |
| EC20111174 | 1 | 2 | 3 | 4 | 5 | 6 | 7 | 8 | 9 |
| EC20111175 | 1 | 2 | 3 | 4 | 5 | 6 | 7 | 8 | 9 |
| EC20120002 | 1 | 2 | 3 | 4 | 5 | 6 | 7 | 8 | 9 |
| EC20120005 | 1 | 2 | 3 | 4 | 5 | 6 | 7 | 8 | 9 |
| EC20120008 | 1 | 2 | 3 | 4 | 5 | 6 | 7 | 8 | 9 |
| EC20121175 | 1 | 2 | 3 | 4 | 5 | 6 | 7 | 8 | 9 |
| P125109 | 1 | | 2 | 3 | 4 | 5 | 6 | 7 | 8 |
| ENTERITIDIS | A | B | C | D | E | F | G | H | I |

| | | | | | | | | |
|---|---|---|---|---|---|---|---|---|
| USDA-ARS-USMARC-1175 | | | | | | | 1 | 2 |
| USDA-ARS-USMARC-1676 | | | | | | | 1 | 2 |
| USDA-ARS-USMARC-1677 | | | | | | | 1 | 2 |
| USDA-ARS-USMARC-1727 | | | | | | | 1 | 2 |
| USDA-ARS-USMARC-1728 | | | | | | | 1 | 2 |
| USDA-ARS-USMARC-1735 | 1 | 2 | 3 | 4 | 5 | 6 | 7 | 8 |
| USDA-ARS-USMARC-1736 | 1 | 2 | 3 | 4 | 5 | 6 | 7 | 8 |
| USDA-ARS-USMARC-1765 | | | | | | | 1 | 2 |
| USDA-ARS-USMARC-1766 | 1 | 2 | 3 | 4 | 5 | 6 | 7 | 8 |
| USDA-ARS-USMARC-1781 | 1 | 2 | 3 | 4 | 5 | 6 | 7 | 8 |
| USDA-ARS-USMARC-1783 | | | | | | | 1 | 2 |
| ATCC BAA-1592 | 1 | 2 | 3 | 4 | 5 | 6 | 7 | 8 |
| CDC 06-0532 | 1 | 2 | 3 | 4 | 5 | 6 | 7 | 8 |
| ANATUM | A | B | C | D | E | F | G | H |

| | | | | | | | |
|---|---|---|---|---|---|---|---|
| AKU_12601 | 1 | 2 | 3 | 4 | 5 | 6 | 7 |
| ATCC 11511 | 1 | 2 | 3 | 4 | 5 | 6 | 7 |
| ATCC 9150 | 1 | 2 | 3 | | | 4 | 5 |
| *PARATYPHI A | A | B | C | D | E | F | G |

| | | | |
|---|---|---|---|
| ATCC 9120 | 1 | 2 | |
| S06004 | | 1 | 2 |
| *PULLORUM | A | B | C |

| | | |
|---|---|---|
| CDC1983-67 | 1 | 2 |
| RKS5078 | 1 | 2 |
| *GALLINARIUM/PULLORUM | A | B |

| | | |
|---|---|---|
| 287/91 | 1 | 2 |
| 9184 | 1 | 2 |
| * GALLINARIUM | A | B |

| | | |
|---|---|---|
| CT_02021853 | 1 | 2 |
| DUBLIN | A | B |

| | | | | | | | | | | | | | | | | | | | | | | | | | | | |
|---|---|---|---|---|---|---|---|---|---|---|---|---|---|---|---|---|---|---|---|---|---|---|---|---|---|---|---|
| CVM 21538 | 1 | 2 | 3 | 4 | 5 | 6 | 7 | 8 | 9 | 10 | 11 | 12 | 13 | 14 | 15 | 16 | 17 | 18 | 19 | | 20 | 21 | 22 | 23 | 24 | 25 | 26 |
| CVM 21550 | 1 | 2 | 3 | 4 | 5 | 6 | 7 | 8 | 9 | 10 | 11 | 12 | 13 | 14 | 15 | 16 | 17 | 18 | 19 | | 20 | 21 | 22 | 23 | 24 | 25 | 26 |
| CVM 22425 | 1 | 2 | 3 | 4 | 5 | 6 | 7 | 8 | 9 | 10 | 11 | 12 | 13 | 14 | 15 | 16 | 17 | 18 | 19 | | 20 | 21 | 22 | 23 | 24 | 25 | 26 |
| CVM 22462 | 1 | 2 | 3 | 4 | 5 | 6 | 7 | 8 | 9 | 10 | 11 | 12 | 13 | 14 | 15 | 16 | 17 | 18 | 19 | | 20 | 21 | 22 | 23 | 24 | 25 | 26 |
| CVM 22513 | 1 | 2 | 3 | 4 | 5 | 6 | 7 | 8 | 9 | 10 | 11 | 12 | 13 | | | | | | | | 14 | 15 | 16 | 17 | 18 | 19 | 20 |
| CVM N1543 | 1 | 2 | 3 | 4 | 5 | 6 | 7 | 8 | 9 | 10 | 11 | 12 | 13 | 14 | 15 | 16 | 17 | 18 | 19 | | 20 | 21 | 22 | 23 | 24 | 25 | 26 |
| CVM N18486 | 1 | 2 | 3 | 4 | 5 | 6 | 7 | | | | | | | | | | | 8 | 9 | 10 | 11 | 12 | 13 | 14 | 15 | 16 | 17 |
| SL254 | 1 | 2 | 3 | 4 | 5 | 6 | 7 | 8 | 9 | 10 | 11 | 12 | 13 | 14 | 15 | 16 | 17 | 18 | 19 | | 20 | 21 | 22 | 23 | 24 | 25 | 26 |
| USMARC-S3124.1 | 1 | 2 | 3 | 4 | 5 | 6 | 7 | 8 | 9 | 10 | 11 | 12 | 13 | 14 | 15 | 16 | 17 | 18 | 19 | | 20 | 21 | 22 | 23 | 24 | 25 | 26 |
| WA_14882 | 1 | 2 | 3 | 4 | 5 | 6 | 7 | 8 | 9 | 10 | 11 | 12 | | 13 | | 14 | 15 | 16 | 17 | 18 | 19 | 20 | 21 | 22 | 23 | 24 | 25 |
| NEWPORT II | A | B | C | D | E | F | G | H | I | J | K | L | M | N | O | P | Q | R | S | T | U | V | W | X | Y | Z | AA |

| | | | | | | | |
|---|---|---|---|---|---|---|---|
| P-stx-12 | 1 | | 2 | 3 | 4 | 5 | 6 |
| Ty2 | 1 | | 2 | 3 | 4 | 5 | 6 |
| CT18 | 1 | 2 | | 3 | 4 | 5 | 6 |
| 1016889 | 1 | | 2 | 3 | 4 | 5 | 6 |
| 1036491 | 1 | | 2 | 3 | 4 | 5 | 6 |
| 129-0238-M | 1 | | 2 | 3 | 4 | 5 | 6 |
| ERL034151 | 1 | 2 | 3 | 4 | 5 | 6 | 7 |
| * TYPHI | A | | B | C | D | E | F |

| | | | | | | | | | | | | | | | | | | | | | | | | |
|---|---|---|---|---|---|---|---|---|---|---|---|---|---|---|---|---|---|---|---|---|---|---|---|---|
| 3114 | 1 | 2 | 3 | 4 | 5 | 6 | 7 | 8 | 9 | 10 | 11 | 12 | 13 | 14 | 15 | 16 | 17 | 18 | 19 | 20 | 21 | 22 | 23 | 24 |
| BOVISMORBIFICANS | A | B | C | D | E | F | G | H | I | J | K | L | M | N | O | P | Q | R | S | T | U | V | W | X |

| | | | | | | | | | | | | | | | | | | | | | | | | | | | | | | | | |
|---|---|---|---|---|---|---|---|---|---|---|---|---|---|---|---|---|---|---|---|---|---|---|---|---|---|---|---|---|---|---|---|---|
| ATCC 10722 | 1 | 2 | 3 | 4 | 5 | 6 | 7 | 8 | 9 | 10 | 11 | 12 | 13 | 14 | 15 | 16 | 17 | 18 | 19 | 20 | 21 | 22 | 23 | 24 | 25 | 26 | 27 | 28 | 29 | 30 | 31 | 32 |
| CFSAN001387 | 1 | 2 | 3 | 4 | 5 | 6 | 7 | 8 | 9 | 10 | 11 | 12 | 13 | 14 | 15 | 16 | 17 | 18 | 19 | 20 | 21 | 22 | | 23 | | | 25 | | | | | |
| CFSAN070643 | 1 | 2 | 3 | 4 | 5 | 6 | 7 | 8 | 9 | 10 | 11 | 12 | 13 | 14 | 15 | 16 | 17 | 18 | 19 | 20 | 21 | 22 | 23 | 24 | 25 | 26 | 27 | 28 | 29 | 30 | 31 | 32 |
| CFSAN076210 | 1 | 2 | 3 | 4 | 5 | 6 | 7 | 8 | 9 | 10 | 11 | 12 | 13 | 14 | 15 | 16 | 17 | 18 | 19 | 20 | 21 | 22 | | 23 | | | 25 | | | | | |
| PIR00537 | 1 | 2 | 3 | 4 | 5 | 6 | 7 | 8 | 9 | 10 | 11 | 12 | 13 | 14 | 15 | 16 | 17 | 18 | 19 | 20 | 21 | 22 | | 23 | | | 25 | | | | | |
| CFSAN070645 | 1 | 2 | 3 | 4 | 5 | 6 | 7 | 8 | 9 | 10 | 11 | 12 | 13 | 14 | 15 | 16 | 17 | 18 | 19 | | | 20 | 21 | 22 | 23 | 24 | 25 | 26 | 27 | 28 | 29 | 30 |
| TXSC_TXSC08-19 | 1 | 2 | 3 | 4 | 5 | 6 | 7 | 8 | 9 | | | | | | | 10 | 11 | 12 | 13 | 14 | 15 | 16 | | 17 | | | | | | | | |
| TENNESSEE | A | B | C | D | E | F | G | H | I | J | K | L | M | N | O | P | Q | R | S | T | U | V | W | X | Y | Z | AA | AB | AC | AD | AE | AF |

| | | | | | | | | | | | | | | | | | | | | | | | | | | | | | | | |
|---|---|---|---|---|---|---|---|---|---|---|---|---|---|---|---|---|---|---|---|---|---|---|---|---|---|---|---|---|---|---|---|
| ATCC 10722 | 33 | 34 | 35 | 36 | 37 | 38 | 39 | 40 | 41 | 42 | 43 | 44 | 45 | 46 | 47 | 48 | 49 | 50 | 51 | 52 | 53 | 54 | 55 | 56 | 57 | 58 | 59 | 60 | 61 | 62 | 63 |
| CFSAN001387 | | 24 | | 26 | 27 | 28 | 29 | 30 | 31 | 32 | 33 | 34 | 35 | 36 | 37 | 38 | 39 | 40 | 41 | 42 | 43 | 44 | 45 | | | | | | | 46 | 47 |
| CFSAN070643 | 33 | 34 | 35 | 36 | 37 | 38 | 39 | 40 | 41 | 42 | 43 | 44 | 45 | 46 | 47 | 48 | 49 | 50 | 51 | 52 | 53 | 54 | 55 | 56 | 57 | 58 | 59 | 60 | 61 | 62 | 63 |
| CFSAN076210 | | 24 | | 26 | 27 | 28 | 29 | 30 | 31 | 32 | 33 | 34 | 35 | 36 | 37 | 38 | 39 | 40 | 41 | 42 | 43 | 44 | 45 | | | | | | | 46 | 47 |
| PIR00537 | | 24 | | 26 | 27 | 28 | 29 | 30 | 31 | 32 | 33 | 34 | 35 | 36 | 37 | 38 | 39 | 40 | 41 | 42 | 43 | 44 | 45 | | | | | | | 46 | 47 |
| CFSAN070645 | 31 | 32 | 33 | 34 | 35 | 36 | 37 | 38 | 39 | 40 | 41 | 42 | 43 | | | 44 | 45 | 46 | 47 | 48 | 49 | 50 | 51 | 52 | 53 | 54 | 55 | 56 | 57 | 58 | 59 |
| TXSC_TXSC08-19 | | 18 | 19 | 20 | 21 | 22 | 23 | 24 | 25 | 26 | 27 | 28 | 29 | 30 | 31 | 32 | 33 | 34 | 35 | 36 | 37 | 38 | 39 | | | | | | | 40 | 41 |
| TENNESSEE | AG | AH | AI | AJ | AK | AL | AM | AN | AO | AP | AQ | AR | AS | AT | AU | AV | AW | AX | AY | AZ | BA | BB | BC | BD | BE | BF | BG | BH | BI | BJ | BK |

| | | | | | | | | | | | | | | | | | | | | | | | | | |
|---|---|---|---|---|---|---|---|---|---|---|---|---|---|---|---|---|---|---|---|---|---|---|---|---|---|
| CFSAN002064 | 1 | 2 | 3 | 4 | 5 | 6 | 7 | 8 | 9 | 10 | 11 | 12 | 13 | 14 | 15 | 16 | 17 | 18 | 19 | 20 | 21 | 21 | 23 | 24 | 25 |
| 41578 | 1 | 2 | 3 | 4 | 5 | 6 | 7 | 8 | 9 | 10 | 11 | 12 | 13 | 14 | 15 | 16 | 17 | 18 | 19 | 20 | 21 | 21 | 23 | 24 | 25 |
| B182 | 1 | 2 | 3 | 4 | 5 | 6 | 7 | 8 | 9 | 10 | 11 | 12 | 13 | 14 | 15 | 16 | 17 | 18 | 19 | 20 | 21 | 22 | 23 | 24 | 25 |
| CFSAN002069 | 1 | 2 | 3 | 4 | 5 | 6 | 7 | 8 | 9 | 10 | 11 | 12 | 13 | 14 | 15 | 16 | 17 | 18 | 19 | 20 | 21 | 22 | 23 | 24 | 25 |
| SARA35 | 1 | 2 | 3 | 4 | 5 | 6 | 7 | 8 | 9 | 10 | 11 | 12 | 13 | 14 | 15 | 16 | 17 | 18 | 19 | 20 | 21 | 22 | 23 | 24 | 25 |
| SL476 | 1 | 2 | 3 | 4 | 5 | 6 | 7 | 8 | 9 | 10 | 11 | 12 | 13 | 14 | 15 | 16 | 17 | 18 | 19 | 20 | 21 | 22 | 23 | 24 | 25 |
| HEIDELBERG | A | B | C | D | E | F | G | H | I | J | K | L | M | N | O | P | Q | R | S | T | U | V | W | X | Y |

| | | | | | | | | | | | | | | | | | | | | | | | | | | | | | | | | | | | | | | | | |
|---|---|---|---|---|---|---|---|---|---|---|---|---|---|---|---|---|---|---|---|---|---|---|---|---|---|---|---|---|---|---|---|---|---|---|---|---|---|---|---|---|
| CFSAN004173 | 1 | 2 | 3 | 4 | 5 | 6 | 7 | 8 | 9 | 10 | 11 | 12 | 13 | 14 | 15 | | | | | | | | | | | | | | | | | | | | | | | | | |
| CFSAN004174 | 1 | 2 | 3 | 4 | 5 | 6 | 7 | 8 | 9 | 10 | 11 | 12 | 13 | 14 | 15 | | | | | | | | | | | | | | | | | | | | | | | | | |
| CFSAN004175 | 1 | 2 | 3 | 4 | 5 | 6 | 7 | 8 | 9 | 10 | 11 | 12 | 13 | 14 | 15 | | | | | | | | | | | | | | | | | | | | | | | | | |
| SGB23 | 1 | 2 | 3 | 4 | 5 | 6 | | | | | 7 | 8 | 9 | 10 | 11 | | | | | | | | | | | | | | | | | | | | | | | | | |
| SARA26 | 1 | 2 | 3 | 4 | 5 | 6 | 7 | 8 | 9 | 10 | 11 | 12 | 13 | 14 | 15 | 16 | 17 | 18 | 19 | 20 | 21 | 22 | 23 | 24 | 25 | 26 | 27 | 28 | 29 | 30 | 31 | 32 | 33 | 35 | 36 | 37 | 38 | 39 | 40 | 41 |
| SAINTPAUL | A | B | C | D | E | F | G | H | I | J | K | L | M | N | O | P | Q | R | S | T | U | V | W | X | Y | Z | AA | AB | AC | AD | AE | AF | AG | AH | AI | AJ | AK | AL | AM | AN |

| | A | B | C | D | E | F | G | H | I | J | K | L | M | N | O | P | Q | R | S | T | U | V | W | X | Y | Z | AA | AB | AC |
|---|---|---|---|---|---|---|---|---|---|---|---|---|---|---|---|---|---|---|---|---|---|---|---|---|---|---|---|---|---|
| USDA-ARS-USMARC-1808 | | 1 | | | | | | | | | | | | | | | | | | | 2 | 3 | 4 | 5 | 6 | 7 | 8 | 9 | 10 |
| USDA-ARS-USMARC-1810 | | 1 | | | | | | 2 | | | 3 | 4 | 5 | 6 | 7 | 8 | 9 | 10 | 11 | 12 | 13 | 14 | 15 | 16 | 17 | 18 | 19 | 20 | 21 |
| USDA-ARS-USMARC-1880 | | 1 | 2 | | | | | | | | | | | | | | | 3 | 4 | 5 | 6 | 7 | 8 | | | | 9 | 10 | 11 |
| USDA-ARS-USMARC-1896 | 1 | 2 | 3 | 4 | 5 | 6 | 7 | 8 | 9 | 10 | 11 | 12 | 13 | 14 | 15 | 16 | | 17 | 18 | 19 | 20 | 21 | 22 | 23 | 24 | 25 | 26 | 27 | 28 |
| USDA-ARS-USMARC-1898 | | 1 | 2 | 3 | | 4 | 5 | 6 | 7 | 8 | 9 | 10 | 11 | 12 | | 13 | 14 | | 15 | 16 | 17 | 18 | 19 | | | | 20 | 21 | 22 |
| USDA-ARS-USMARC-1899 | | 1 | | | | | 2 | 3 | | | 4 | 5 | 6 | | | | | | | | | | | | | | | 7 | 8 |
| str. 14028S | | 1 | | | | | 2 | 3 | | | 4 | 5 | 6 | 7 | 8 | 9 | 10 | 11 | 12 | 13 | 14 | 15 | 16 | 17 | 18 | 19 | 20 | 21 | 22 |
| str. 798 | | 1 | | | | | 2 | 3 | | | 4 | 5 | 6 | 7 | 8 | 9 | 10 | 11 | 12 | 13 | 14 | 15 | 16 | 17 | 18 | 19 | 20 | 21 | 22 |
| CDC 2009K-1640 | | 1 | | | | | | | | | | | | | | | | | | | 2 | 3 | 4 | 5 | 6 | 7 | 8 | 9 | 10 |
| CDC 2009K-2059 | 1 | 2 | 3 | 4 | 5 | 6 | 7 | 8 | 9 | 10 | 11 | 12 | 13 | 14 | 15 | 16 | | 17 | 18 | 19 | 20 | 21 | 22 | 23 | 24 | 25 | 26 | 27 | 28 |
| CDC 2010K-1587 | | 1 | 2 | 3 | 4 | | 5 | 6 | 7 | 8 | 9 | 10 | 11 | 12 | 13 | 14 | 15 | 16 | 17 | 18 | 19 | 20 | 21 | 22 | 23 | 24 | 25 | 26 | 27 |
| CDC 2011K-0870 | | 1 | 2 | | | | | | | | | | | | | | | 3 | 4 | 5 | 6 | 7 | 8 | | | | 9 | 10 | 11 |
| CDC 2011K-1702 | | 1 | | | | | | | | | | | | | | | | | | | 2 | 3 | 4 | 5 | 6 | 7 | 8 | 9 | 10 |
| D23580 | | 1 | | | | | 2 | 3 | | | 4 | 5 | 6 | 7 | 8 | 9 | 10 | 11 | 12 | 13 | 14 | | 15 | 16 | 17 | 18 | 19 | 20 | 21 |
| DT104 | | 1 | | | | | | | | | | | | | | | | | | | 2 | 3 | 4 | 5 | 6 | 7 | 8 | 9 | 10 |
| DT2 | | 1 | | | | | 2 | 3 | | | 4 | 5 | 6 | 7 | 8 | 9 | 10 | 11 | 12 | | 13 | 14 | 15 | 16 | 17 | 18 | | 19 | 20 |
| L-3553 | | 1 | | | | | 2 | 3 | | | 4 | 5 | 6 | 7 | 8 | 9 | 10 | 11 | 12 | 13 | 14 | 15 | 16 | 17 | 18 | 19 | 20 | 21 | 22 |
| SARA13 | | 1 | | | | | 2 | 3 | | | 4 | 5 | 6 | 7 | | | | | | 8 | 9 | 10 | 11 | 12 | 13 | 14 | 15 | 16 | 17 |
| SL1344 | | 1 | | | | | 2 | 3 | | | 4 | 5 | 6 | 7 | | | | | | 8 | | | | 9 | 10 | 11 | 12 | 13 | 14 |
| ST4/74 | | 1 | | | | | 2 | 3 | | | 4 | 5 | 6 | 7 | | | | | | 8 | | | | 9 | 10 | 11 | 12 | 13 | 14 |
| T000240 | | 1 | 2 | 3 | | | | 4 | | 5 | 6 | 7 | 8 | 9 | | 10 | 11 | 12 | 13 | 14 | 15 | 16 | 17 | | | | 18 | 19 | 20 |
| UK-1 | | 1 | | | | | 2 | 3 | | | 4 | 5 | 6 | 7 | | | 8 | 9 | 10 | 11 | 12 | 13 | 14 | | | | | | |
| CFSAN001921 | | 1 | 2 | 3 | 4 | | 5 | 6 | 7 | 8 | 9 | 10 | 11 | 12 | 13 | 14 | 15 | 16 | 17 | 18 | 19 | 20 | 21 | 22 | 23 | 24 | 25 | 26 | 27 |
| TYPHIMURIUM | A | B | C | D | E | F | G | H | I | J | K | L | M | N | O | P | Q | R | S | T | U | V | W | X | Y | Z | AA | AB | AC |

| | | | | | | | | | | | | | | | | | | | | | | | | | | | | | |
|---|---|---|---|---|---|---|---|---|---|---|---|---|---|---|---|---|---|---|---|---|---|---|---|---|---|---|---|---|---|
| 50 MZ0080 | 1 | 2 | 3 | 4 | 5 | 6 | 7 | 8 | 9 | 10 | 11 | 12 | 13 | 14 | 15 | 16 | 17 | 18 | 19 | 20 | 21 | 22 | 23 | 24 | 25 | 26 | 27 | 28 | 29 |
| DIARIZONAE | A | B | C | D | E | F | G | H | I | J | K | L | M | N | O | P | Q | R | S | T | U | V | W | X | Y | Z | AA | AB | AC |

| | | | | | | | | | | | | | | | | | | | | | | | |
|---|---|---|---|---|---|---|---|---|---|---|---|---|---|---|---|---|---|---|---|---|---|---|---|
| NCTC10047 | 1 | 2 | 3 | 4 | 5 | 6 | 7 | 8 | 9 | 10 | 11 | 12 | 13 | 14 | 15 | 16 | 17 | 18 | 19 | 20 | 21 | 22 | 23 |
| ARIZONAE | A | B | C | D | E | F | G | H | I | J | K | L | M | N | O | P | Q | R | S | T | U | V | W |

| | A | B | C | D | E | F | G | H | I | J | K | L | M | N | O | P | Q | R | S | T | U | V | W | X | Y | Z | AA | AB | AC | AD | AE | AF | AG | AH | AI | AJ |
|---|---|---|---|---|---|---|---|---|---|---|---|---|---|---|---|---|---|---|---|---|---|---|---|---|---|---|---|---|---|---|---|---|---|---|---|---|
| FCC0123 | 1 | | | | | | | | | | | | | | | | | | | | | | | | | | | | | | | | | 2 | 3 | 4 |
| CDC 2009K-0792 | | 1 | 2 | 3 | 4 | 5 | 6 | 7 | 8 | 9 | 10 | 11 | 12 | 13 | 14 | 15 | 16 | 17 | 18 | 19 | 20 | 21 | 22 | 23 | 24 | 25 | 26 | 27 | 28 | 29 | 30 | 31 | 32 | 33 | 34 | 35 |
| USDA-ARS-USMARC-1900 | 1 | 2 | 3 | 4 | 5 | 6 | 7 | 8 | 9 | 10 | 11 | 12 | 13 | 14 | 15 | 16 | 17 | 18 | 19 | 20 | 21 | 22 | 23 | 24 | 25 | 26 | 27 | 28 | 29 | 30 | 31 | 32 | 33 | 34 | 35 | 36 |
| USDA-ARS-USMARC-1901 | | 1 | 2 | 3 | 4 | 5 | 6 | 7 | 8 | 9 | 10 | 11 | 12 | 13 | 14 | 15 | 16 | 17 | 18 | 19 | 20 | 21 | 22 | 23 | 24 | | 25 | 26 | 27 | 28 | 29 | 30 | 31 | 32 | 33 | 34 |
| USDA-ARS-USMARC-1904 | | 1 | 2 | 3 | 4 | 5 | 6 | 7 | 8 | 9 | 10 | 11 | 12 | 13 | 14 | 15 | 16 | 17 | 18 | 19 | 20 | | | 21 | 22 | 23 | 24 | 25 | 26 | 27 | 28 | 29 | 30 | 31 | 32 | 33 |
| USDA-ARS-USMARC-1912 | | 1 | 2 | 3 | 4 | 5 | 6 | 7 | 8 | 9 | 10 | 11 | 12 | 13 | 14 | 15 | 16 | 17 | 18 | 19 | 20 | 21 | 22 | 23 | 24 | 25 | 26 | 27 | 28 | 29 | 30 | 31 | 32 | 33 | 34 | 35 |
| USDA-ARS-USMARC-1903 | | 1 | 2 | 3 | 4 | 5 | 6 | 7 | 8 | 9 | 10 | 11 | 12 | 13 | 14 | 15 | 16 | 17 | 18 | 19 | 20 | 21 | 22 | 23 | 24 | 25 | 26 | 27 | 28 | 29 | 30 | 31 | 32 | 33 | 34 | 35 |
| USDA-ARS-USMARC-1921 | | 1 | 2 | 3 | 4 | 5 | 6 | 7 | 8 | 9 | 10 | 11 | 12 | 13 | 14 | 15 | 16 | 17 | 18 | 19 | 20 | 21 | 22 | 23 | 24 | 25 | 26 | 27 | 28 | 29 | 30 | 31 | 32 | 33 | 34 | 35 |
| MONTEVIDEO | A | B | C | D | E | F | G | H | I | J | K | L | M | N | O | P | Q | R | S | T | U | V | W | X | Y | Z | AA | AB | AC | AD | AE | AF | AG | AH | AI | AJ |

| | A | B | C | D | E | F | G | H | I | J | K | L | M | N | O | P | Q | R | S | T | U | V | W | X | Y |
|---|---|---|---|---|---|---|---|---|---|---|---|---|---|---|---|---|---|---|---|---|---|---|---|---|---|
| CFSAN051296 | | 1 | 2 | 3 | 4 | 5 | 6 | 7 | 8 | 9 | 10 | 11 | 12 | 13 | 14 | 15 | 16 | 17 | 18 | 19 | 20 | 21 | 22 | 23 | 24 |
| 531954 | | 1 | 2 | 3 | 4 | 5 | 6 | 7 | 8 | 9 | 10 | 11 | 12 | 13 | 14 | 15 | 16 | 17 | 18 | 19 | 20 | 21 | 22 | 23 | 24 |
| 507440-20 | | 1 | 2 | 3 | 4 | 5 | 6 | 7 | 8 | 9 | 10 | 11 | 12 | 13 | 14 | 15 | 16 | 17 | 18 | 19 | 20 | 21 | 22 | 23 | 24 |
| CDC 07-0954 | 1 | 2 | 3 | 4 | 5 | | | | | | | | | | | | | | | 6 | 7 | 8 | 9 | 10 | 11 |
| CDC 08-1942 | | 1 | | | | | | | | | | | | | | | | | | | 2 | 3 | 4 | 5 | 6 |
| CDC 2010K-0257 | | 1 | 2 | 3 | 4 | 5 | 6 | 7 | 8 | 9 | 10 | 11 | 12 | 13 | 14 | 15 | 16 | 17 | 18 | 19 | 20 | 21 | 22 | 23 | 24 |
| CDC 2011K-1674 | 1 | 2 | 3 | 4 | 5 | | | | | | | | | | | | | | | 6 | 7 | 8 | 9 | 10 | 11 |
| CDC 2012K-1544 | | 1 | 2 | 3 | 4 | 5 | 6 | 7 | 8 | 9 | 10 | 11 | 12 | 13 | 14 | 15 | 16 | 17 | 18 | 19 | 20 | 21 | 22 | 23 | 24 |
| CDC 2013K-0218 | | 1 | | | | | | | | | | | | | | | | | | | 2 | 3 | 4 | 5 | 6 |
| MONTEVIDEO | A | B | C | D | E | F | G | H | I | J | K | L | M | N | O | P | Q | R | S | T | U | V | W | X | Y |

| | | | | | | | | | | | | | | | | | | | | |
|---|---|---|---|---|---|---|---|---|---|---|---|---|---|---|---|---|---|---|---|---|
| 66 SA19983605 | 1 | 2 | 3 | 4 | 5 | 6 | 7 | 8 | 9 | 10 | 11 | 12 | 13 | 14 | 15 | 16 | 17 | 18 | 19 | 20 |
| BONGORI | A | B | C | D | E | F | G | H | I | J | K | L | M | N | O | P | Q | R | S | T |

| | A | B | C | D | E | F | G | H | I | J | K | L | M | N | O | P | Q | R |
|---|---|---|---|---|---|---|---|---|---|---|---|---|---|---|---|---|---|---|
| CFSAN001660 | 1 | 2 | 3 | 4 | 5 | 6 | 7 | | | | 8 | 9 | 10 | 11 | 12 | 13 | 14 | 15 |
| CDC 2009K-1331 | 1 | 2 | 3 | | | | | | 4 | 5 | 6 | 7 | 8 | 9 | 10 | 11 | 12 | 13 |
| CDC 2012K-0938 | 1 | | | | | | | | | | | | | | | 2 | 3 | 4 |
| Levine 1 | 1 | 2 | 3 | | | | | | 4 | 5 | 6 | 7 | 8 | 9 | 10 | 11 | 12 | 13 |
| USDA-ARS-USMARC-1927 | 1 | 2 | 3 | 4 | 5 | 6 | 7 | 8 | 9 | 10 | 11 | 12 | 13 | 14 | 15 | 16 | 17 | 18 |
| Levine 15 | 1 | | | | | | | | | | | | | | 2 | 3 | 4 | 5 |
| NEWPORT III | A | B | C | D | E | F | G | H | I | J | K | L | M | N | O | P | Q | R |

**B**

| | | | | | | | | | | | | | | | | | | | | | | | | | | |
|---|---|---|---|---|---|---|---|---|---|---|---|---|---|---|---|---|---|---|---|---|---|---|---|---|---|---|
| USDA-ARS-USMARC-1175 | 1 | 2 | 3 | 4 | 5 | 6 | 7 | 8 | 9 | 10 | 11 | 12 | 13 | 14 | 15 | 16 | 17 | 18 | 19 | 20 | 21 | 22 | 23 | 24 | 25 | 26 |
| USDA-ARS-USMARC-1676 | 1 | 2 | | | | | | | | 3 | 4 | 5 | 6 | 7 | 8 | 9 | 10 | 11 | 12 | 13 | 14 | 15 | 16 | 17 | 18 | 19 |
| USDA-ARS-USMARC-1677 | | | | | | 1 | 2 | 3 | 4 | 5 | 6 | 7 | 8 | 9 | 10 | 11 | 12 | 13 | 14 | 15 | 16 | 17 | 18 | 19 | 20 | 21 |
| USDA-ARS-USMARC-1727 | 1 | 2 | 3 | | | | | | | | | | | | | | | 4 | 5 | 6 | 7 | 8 | 9 | 10 | 11 | 12 |
| USDA-ARS-USMARC-1728 | 1 | 2 | 3 | 4 | 5 | 6 | 7 | 8 | 9 | | 10 | 11 | 12 | 13 | 14 | 15 | 16 | 17 | 18 | 19 | 20 | 21 | 22 | 23 | 24 | 25 |
| USDA-ARS-USMARC-1735 | | | | | | | | | | | | | | | | | | 1 | 2 | | 3 | 4 | 5 | 6 | 7 | 8 |
| USDA-ARS-USMARC-1736 | 1 | 2 | 3 | 4 | 5 | 6 | 7 | 8 | 9 | 10 | 11 | 12 | 13 | 14 | 15 | 16 | 17 | 18 | 19 | 20 | 21 | 22 | 23 | 24 | 25 | 26 |
| USDA-ARS-USMARC-1765 | 1 | 2 | 3 | 4 | 5 | 6 | | | 7 | 8 | 9 | 10 | 11 | | | | 12 | 13 | 14 | 15 | 16 | 17 | 18 | 19 | 20 | 21 |
| USDA-ARS-USMARC-1766 | 1 | 2 | 3 | 4 | 5 | 6 | 7 | 8 | 9 | 10 | 11 | 12 | 13 | 14 | 15 | 16 | 17 | 18 | 19 | 20 | 21 | 22 | 23 | 24 | | |
| USDA-ARS-USMARC-1781 | 1 | 2 | 3 | 4 | 5 | 6 | 7 | 8 | 9 | 10 | | | | | | | 11 | 12 | 13 | 14 | 15 | 16 | 17 | 18 | 19 | 20 |
| USDA-ARS-USMARC-1783 | 1 | 2 | 3 | 4 | 5 | 6 | 7 | 8 | 9 | | 10 | 11 | 12 | 13 | 14 | 15 | 16 | 17 | 18 | 19 | 20 | 21 | 22 | 23 | 24 | 25 |
| ATCC BAA-1592 | 1 | 2 | 3 | 4 | 5 | 6 | 7 | 8 | 9 | 10 | | | | | | | 11 | 12 | 13 | 14 | 15 | 16 | 17 | 18 | 19 | 20 |
| CDC 06-0532 | 1 | 2 | 3 | 4 | 5 | 6 | 7 | 8 | 9 | 10 | | | | | | | 11 | 12 | 13 | 14 | 15 | 16 | 17 | 18 | 19 | 20 |
| ANATUM | A | B | C | D | E | F | G | H | I | J | K | L | M | N | O | P | Q | R | S | T | U | V | W | X | Y | Z |

| | | | | | | | | | | | | |
|---|---|---|---|---|---|---|---|---|---|---|---|---|
| 77-1427 | 1 | | | | | 2 | 3 | 4 | 5 | 6 | 7 | 8 |
| CDC_2010K_0968 | 1 | 2 | 3 | | 4 | 5 | 6 | 7 | 8 | 9 | 10 | 11 |
| EC20090641 | 1 | 2 | 3 | | 4 | 5 | 6 | 7 | 8 | 9 | 10 | 11 |
| EC20090698 | 1 | 2 | 3 | | 4 | 5 | 6 | 7 | 8 | 9 | 10 | 11 |
| EC20110221 | 1 | 2 | 3 | | 4 | 5 | 6 | 7 | 8 | 9 | 10 | 11 |
| EC20110354 | 1 | 2 | 3 | 4 | 5 | 6 | 7 | 8 | 9 | 10 | 11 | 12 |
| EC20110355 | 1 | 2 | 3 | 4 | 5 | 6 | 7 | 8 | 9 | 10 | 11 | 12 |
| EC20110356 | 1 | 2 | 3 | 4 | 5 | 6 | 7 | 8 | 9 | 10 | 11 | 12 |
| EC20110357 | 1 | 2 | 3 | | 4 | 5 | 6 | 7 | 8 | 9 | 10 | 11 |
| EC20110358 | 1 | 2 | 3 | | 4 | 5 | 6 | 7 | 8 | 9 | 10 | 11 |
| EC20110360 | 1 | 2 | 3 | 4 | 5 | 6 | 7 | 8 | 9 | 10 | 11 | 12 |
| EC20110361 | 1 | 2 | 3 | 4 | 5 | 6 | 7 | 8 | 9 | 10 | 11 | 12 |
| EC20111095 | 1 | 2 | 3 | 4 | 5 | 6 | 7 | 8 | 9 | 10 | 11 | 12 |
| EC20111174 | 1 | 2 | 3 | 4 | 5 | 6 | 7 | 8 | 9 | 10 | 11 | 12 |
| EC20111175 | 1 | 2 | 3 | 4 | 5 | 6 | 7 | 8 | 9 | 10 | 11 | 12 |
| EC20120002 | 1 | 2 | 3 | 4 | 5 | 6 | 7 | 8 | 9 | 10 | 11 | 12 |
| EC20120005 | 1 | 2 | 3 | 4 | 5 | 6 | 7 | 8 | 9 | 10 | 11 | 12 |
| EC20120008 | 1 | 2 | 3 | | 4 | 5 | 6 | 7 | 8 | 9 | 10 | 11 |
| EC20121175 | 1 | 2 | 3 | 4 | 5 | 6 | 7 | 8 | 9 | 10 | 11 | 12 |
| P125109 | 1 | 2 | | | 3 | 4 | 5 | 6 | 7 | 8 | 9 | 10 |
| ENTERITIDIS | A | B | C | D | E | F | G | H | I | J | K | L |

| | | | | | | | | | | | | | | | | | | |
|---|---|---|---|---|---|---|---|---|---|---|---|---|---|---|---|---|---|---|
| CFSAN002064 | 1 | 2 | 3 | 4 | 5 | 6 | 7 | 8 | 9 | 10 | 11 | 12 | 13 | 14 | 15 | 16 | 17 | 18 |
| 41578 | 1 | 2 | 3 | 4 | 5 | | | 6 | 7 | 8 | 9 | 10 | 11 | 12 | 13 | 14 | 15 | 16 |
| B182 | 1 | 2 | 3 | 4 | 5 | 6 | 7 | 8 | 9 | 10 | 11 | 12 | 13 | 14 | 15 | 16 | 17 | 18 |
| CFSAN002069 | 1 | 2 | 3 | 4 | 5 | 6 | 7 | 8 | 9 | 10 | 11 | 12 | 13 | 14 | 15 | 16 | 17 | 18 |
| SARA35 | 1 | 2 | 3 | 4 | 5 | 6 | 7 | 8 | 9 | 10 | 11 | 12 | 13 | 14 | 15 | 16 | 17 | 18 |
| SL476 | 1 | 2 | 3 | 4 | 5 | 6 | 7 | 8 | 9 | 10 | 11 | 12 | 13 | 14 | 15 | 16 | 17 | 18 |
| HEIDELBERG | A | B | C | D | E | F | G | H | I | J | K | L | M | N | O | P | Q | R |

| | | | |
|---|---|---|---|
| AKU_12601 | 1 | 2 | 3 |
| ATCC 11511 | 1 | 2 | 3 |
| ATCC 9150 | 1 | 2 | 3 |
| *PARATYPHI A | A | B | C |

| | |
|---|---|
| P-stx-12 | 1 |
| Ty2 | 1 |
| CT18 | 1 |
| 1016889 | 1 |
| 1036491 | 1 |
| 129-0238-M | 1 |
| ERL034151 | 1 |
| * TYPHI | A |

| | | | | | | |
|---|---|---|---|---|---|---|
| ATCC 9120 | 1 | 2 | 3 | 4 | 5 | 6 |
| S06004 | 1 | | | | | 2 |
| *PULLORUM | A | B | C | D | E | F |

| | | | | | | | | | | |
|---|---|---|---|---|---|---|---|---|---|---|
| 287/91 | 1 | 2 | 3 | 4 | 5 | 6 | 7 | 8 | 9 | 10 |
| 9184 | 1 | 2 | 3 | 4 | 5 | 6 | 7 | 8 | 9 | 10 |
| *GALLINARIUM | A | B | C | D | E | F | G | H | I | J |

| | | | | | | |
|---|---|---|---|---|---|---|
| CDC1983-67 | 1 | 2 | 3 | 4 | 5 | 6 |
| RKS5078 | 1 | 2 | 3 | 4 | 5 | 6 |
| *GALLINARIUM/PULLORUM | A | B | C | D | E | F |

| | | | | | | | | | | | | | | | | | | | | | | |
|---|---|---|---|---|---|---|---|---|---|---|---|---|---|---|---|---|---|---|---|---|---|---|
| CFSAN004173 | 1 | 2 | 3 | 4 | 5 | 6 | 7 | 8 | 9 | 10 | | | 11 | 12 | 13 | 14 | 15 | 16 | 17 | 18 | 19 | 20 |
| CFSAN004174 | 1 | 2 | 3 | 4 | 5 | 6 | 7 | 8 | 9 | 10 | | | 11 | 12 | 13 | 14 | 15 | 16 | 17 | 18 | 19 | 20 |
| CFSAN004175 | 1 | 2 | 3 | 4 | 5 | 6 | 7 | 8 | 9 | 10 | | | 11 | 12 | 13 | 14 | 15 | 16 | 17 | 18 | 19 | 20 |
| SGB23 | 1 | 2 | 3 | 4 | 5 | 6 | 7 | 8 | 9 | 10 | 11 | 12 | 13 | 14 | 15 | 16 | 17 | 16 | 19 | 20 | 21 | 22 |
| SARA26 | 1 | 2 | 3 | 4 | 5 | 6 | 7 | 8 | 9 | 10 | 11 | 12 | 13 | 14 | 15 | 16 | 17 | 18 | 19 | 20 | 21 | 22 |
| SAINTPAUL | A | B | C | D | E | F | G | H | I | J | K | L | M | N | O | P | Q | R | S | T | U | V |

| | | | | | |
|---|---|---|---|---|---|
| CT_02021853 | 1 | 2 | 3 | 4 | 5 |
| DUBLIN | A | B | C | D | E |

| | |
|---|---|
| NCTC5772 | 1 |
| *SENDAI | A |

| | | | | | | | | | | | | | | | | | | | | | |
|---|---|---|---|---|---|---|---|---|---|---|---|---|---|---|---|---|---|---|---|---|---|
| CVM 21538 | 1 | 2 | 3 | 4 | 5 | 6 | 7 | 8 | 9 | 10 | 11 | 12 | 13 | 14 | | | 15 | 16 | 17 | 18 | 19 |
| CVM 21550 | 1 | 2 | 3 | 4 | 5 | 6 | 7 | 8 | 9 | 10 | 11 | 12 | 13 | 14 | | | 15 | 16 | 17 | 18 | 19 |
| CVM 22425 | 1 | 2 | 3 | 4 | 5 | 6 | 7 | 8 | 9 | 10 | 11 | 12 | 13 | 14 | | | 15 | 16 | 17 | 18 | 19 |
| CVM 22462 | 1 | 2 | 3 | 4 | 5 | 6 | 7 | 8 | 9 | 10 | 11 | 12 | 13 | 14 | | | 15 | 16 | 17 | 18 | 19 |
| CVM 22513 | 1 | 2 | 3 | 4 | 5 | 6 | 7 | 8 | 9 | 10 | 11 | 12 | 13 | 14 | | | 15 | 16 | 17 | 18 | 19 |
| CVM N1543 | 1 | 2 | 3 | 4 | 5 | 6 | 7 | 8 | 9 | 10 | 11 | 12 | 13 | 14 | | | 15 | 16 | 17 | 18 | 19 |
| CVM N18486 | 1 | 2 | 3 | 4 | 5 | 6 | 7 | 8 | | | | | | | | | | 9 | 10 | 11 | 12 |
| SL254 | 1 | 2 | 3 | 4 | 5 | 6 | 7 | 8 | 9 | 10 | 11 | 12 | 13 | 14 | | | 15 | 16 | 17 | 18 | 19 |
| USMARC-S3124.1 | 1 | 2 | 3 | 4 | 5 | 6 | 7 | 8 | 9 | 10 | 11 | 12 | 13 | 14 | | | 15 | 16 | 17 | 18 | 19 |
| WA_14882 | 1 | 2 | 3 | 4 | 5 | 6 | 7 | 8 | | | 9 | 10 | 11 | 12 | 13 | 14 | 15 | 16 | 17 | 18 | 19 |
| NEWPORT II | A | B | C | D | E | F | G | H | I | J | K | L | M | N | O | P | Q | R | S | T | U |

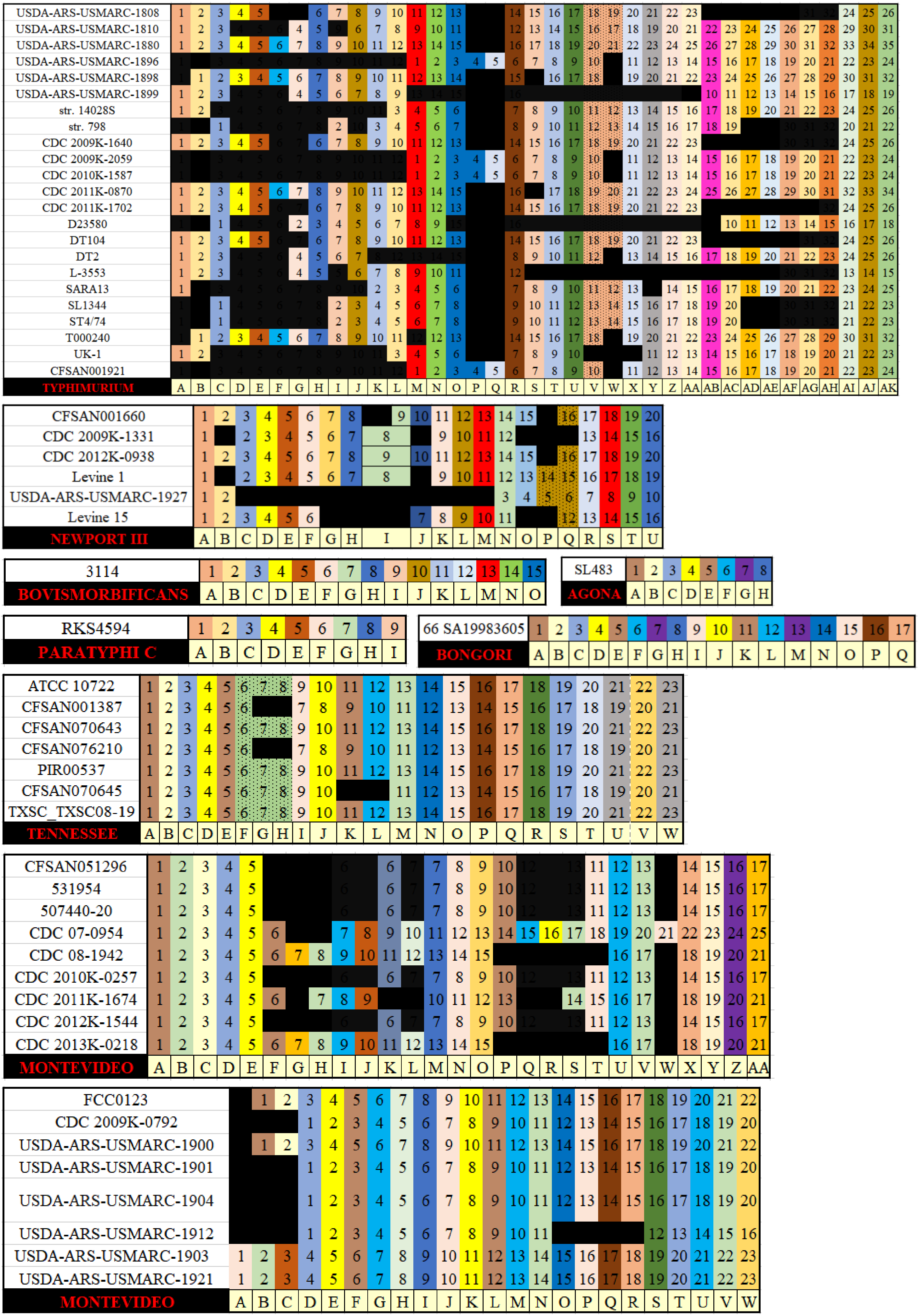
USDA-ARS-USMARC-1808
USDA-ARS-USMARC-1810
USDA-ARS-USMARC-1880
USDA-ARS-USMARC-1896
USDA-ARS-USMARC-1898
USDA-ARS-USMARC-1899
str. 14028S
str. 798
CDC 2009K-1640
CDC 2009K-2059
CDC 2010K-1587
CDC 2011K-0870
CDC 2011K-1702
D23580
DT104
DT2
L-3553
SARA13
SL1344
ST4/74
T000240
UK-1
CFSAN001921
TYPHIMURIUM
CFSAN001660
CDC 2009K-1331
CDC 2012K-0938
Levine 1
USDA-ARS-USMARC-1927
Levine 15
NEWPORT III
3114
BOVISMORBIFICANS
SL483
AGONA
RKS4594
PARATYPHI C
66 SA19983605
BONGORI
ATCC 10722
CFSAN001387
CFSAN070643
CFSAN076210
PIR00537
CFSAN070645
TXSC_TXSC08-19
TENNESSEE
CFSAN051296
531954
507440-20
CDC 07-0954
CDC 08-1942
CDC 2010K-0257
CDC 2011K-1674
CDC 2012K-1544
CDC 2013K-0218
MONTEVIDEO
FCC0123
CDC 2009K-0792
USDA-ARS-USMARC-1900
USDA-ARS-USMARC-1901
USDA-ARS-USMARC-1904
USDA-ARS-USMARC-1912
USDA-ARS-USMARC-1903
USDA-ARS-USMARC-1921
MONTEVIDEO

**C**

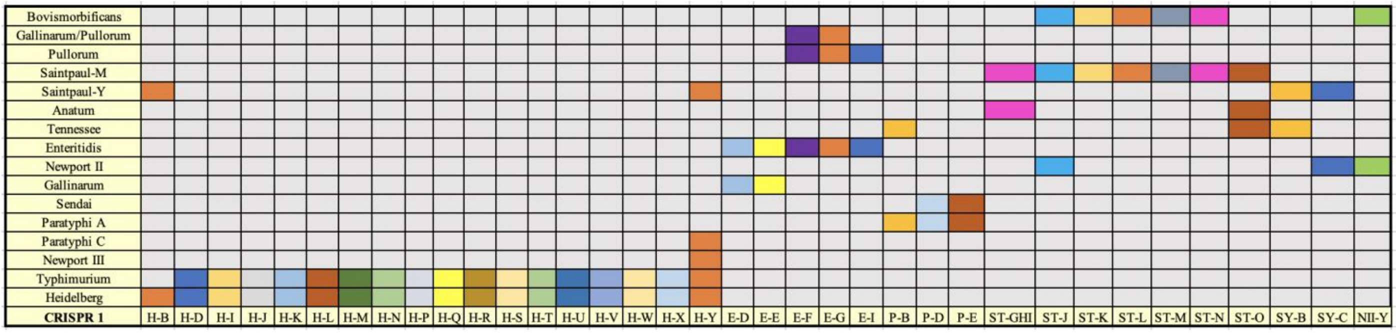
Bovismorbificans
Gallinarum/Pullorum
Pullorum
Saintpaul-M
Saintpaul-Y
Anatum
Tennessee
Enteritidis
Newport II
Gallinarum
Sendai
Paratyphi A
Paratyphi C
Newport III
Typhimurium
Heidelberg
CRISPR 1
H-B H-D H-I H-J H-K H-L H-M H-N H-P H-Q H-R H-S H-T H-U H-V H-W H-X H-Y E-D E-E E-F E-G E-I P-B P-D P-E ST-GHI ST-J ST-K ST-L ST-M ST-N ST-O SY-B SY-C NII-Y

**D**

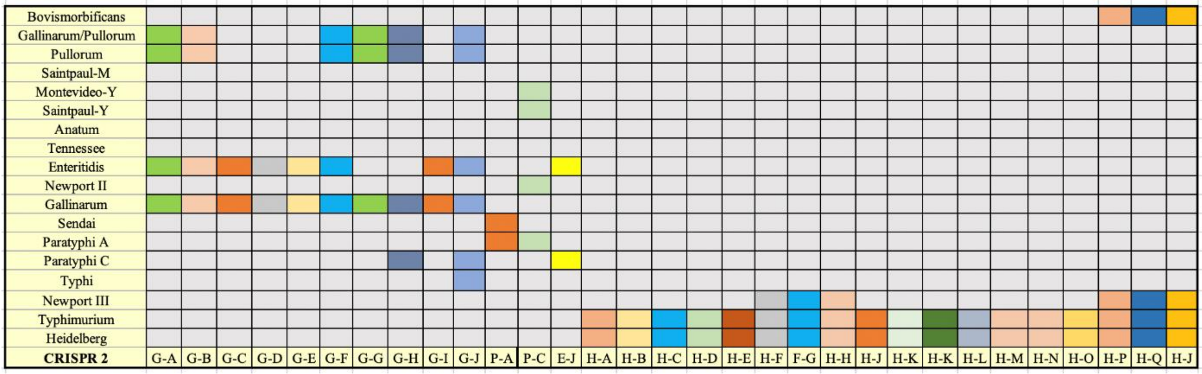
Bovismorbificans
Gallinarum/Pullorum
Pullorum
Saintpaul-M
Montevideo-Y
Saintpaul-Y
Anatum
Tennessee
Enteritidis
Newport II
Gallinarum
Sendai
Paratyphi A
Paratyphi C
Typhi
Newport III
Typhimurium
Heidelberg
CRISPR 2
G-A G-B G-C G-D G-E G-F G-G G-H G-I G-J P-A P-C E-J H-A H-B H-C H-D H-E H-F F-G H-H H-J H-K H-K H-L H-M H-N H-O H-P H-Q H-J

#### Figure S2

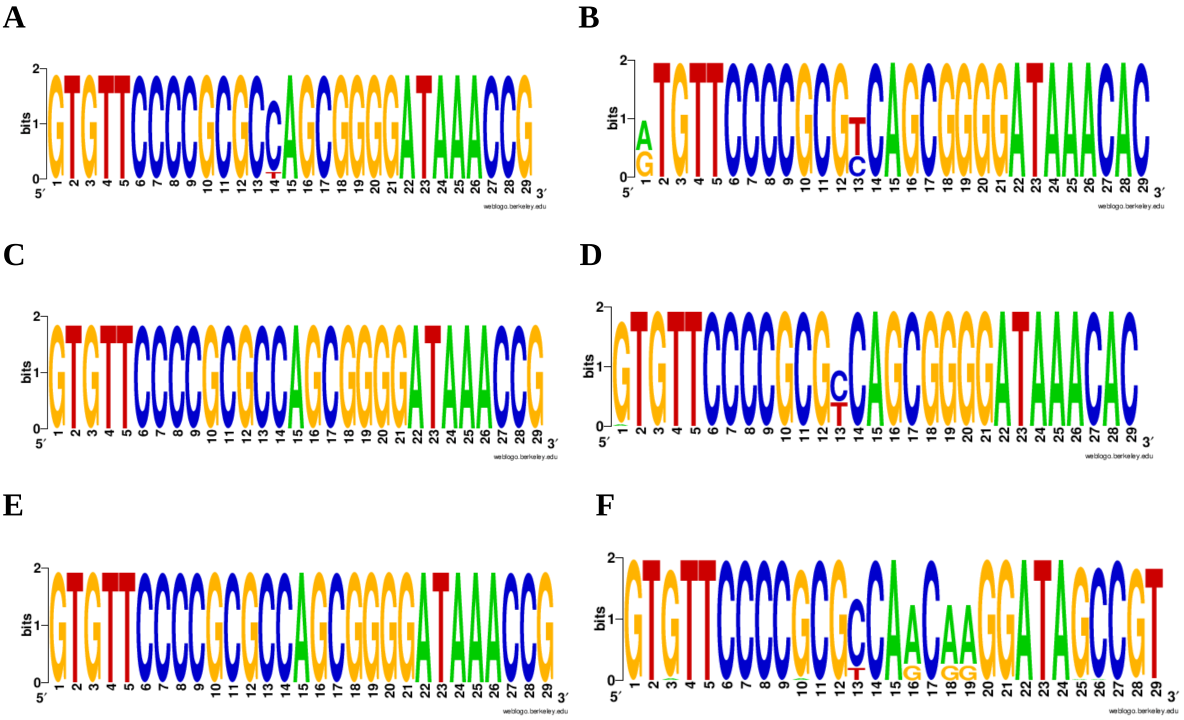
A
B
C
D
E
F
bits
weblogo.berkeley.edu

**Figure S3**

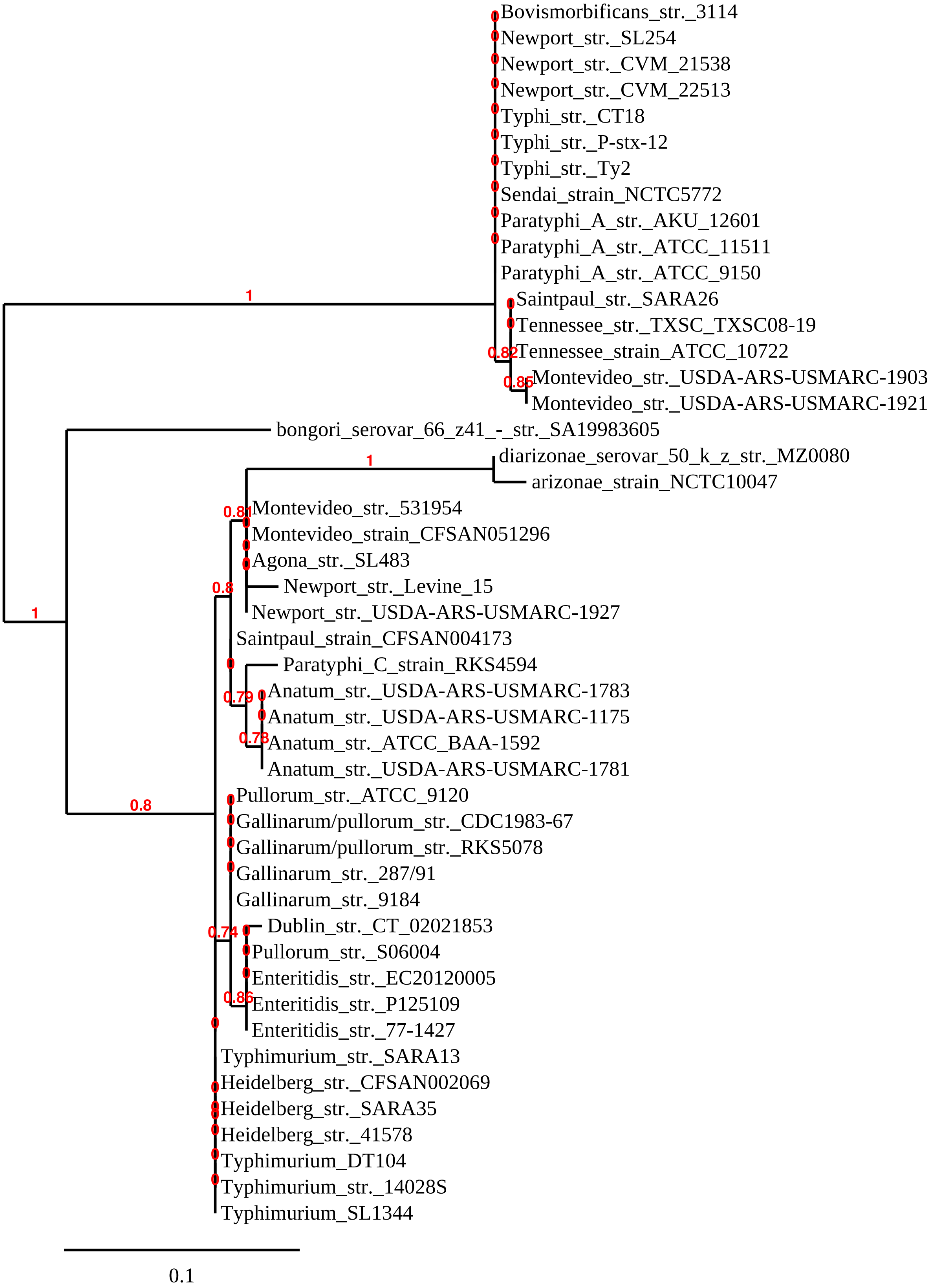
Bovismorbificans_str._3114
Newport_str._SL254
Newport_str._CVM_21538
Newport_str._CVM_22513
Typhi_str._CT18
Typhi_str._P-stx-12
Typhi_str._Ty2
Sendai_strain_NCTC5772
Paratyphi_A_str._AKU_12601
Paratyphi_A_str._ATCC_11511
Paratyphi_A_str._ATCC_9150
Saintpaul_str._SARA26
Tennessee_str._TXSC_TXSC08-19
Tennessee_strain_ATCC_10722
Montevideo_str._USDA-ARS-USMARC-1903
Montevideo_str._USDA-ARS-USMARC-1921
bongori_serovar_66_z41_-_str._SA19983605
diarizonae_serovar_50_k_z_str._MZ0080
arizonae_strain_NCTC10047
Montevideo_str._531954
Montevideo_strain_CFSAN051296
Agona_str._SL483
Newport_str._Levine_15
Newport_str._USDA-ARS-USMARC-1927
Saintpaul_strain_CFSAN004173
Paratyphi_C_strain_RKS4594
Anatum_str._USDA-ARS-USMARC-1783
Anatum_str._USDA-ARS-USMARC-1175
Anatum_str._ATCC_BAA-1592
Anatum_str._USDA-ARS-USMARC-1781
Pullorum_str._ATCC_9120
Gallinarum/pullorum_str._CDC1983-67
Gallinarum/pullorum_str._RKS5078
Gallinarum_str._287/91
Gallinarum_str._9184
Dublin_str._CT_02021853
Pullorum_str._S06004
Enteritidis_str._EC20120005
Enteritidis_str._P125109
Enteritidis_str._77-1427
Typhimurium_str._SARA13
Heidelberg_str._CFSAN002069
Heidelberg_str._SARA35
Heidelberg_str._41578
Typhimurium_DT104
Typhimurium_str._14028S
Typhimurium_SL1344
1
0.82
0.86
1
0.81
0.8
1
0.79
0.78
0.8
0.74
0.86
0.1

**Figure S4**

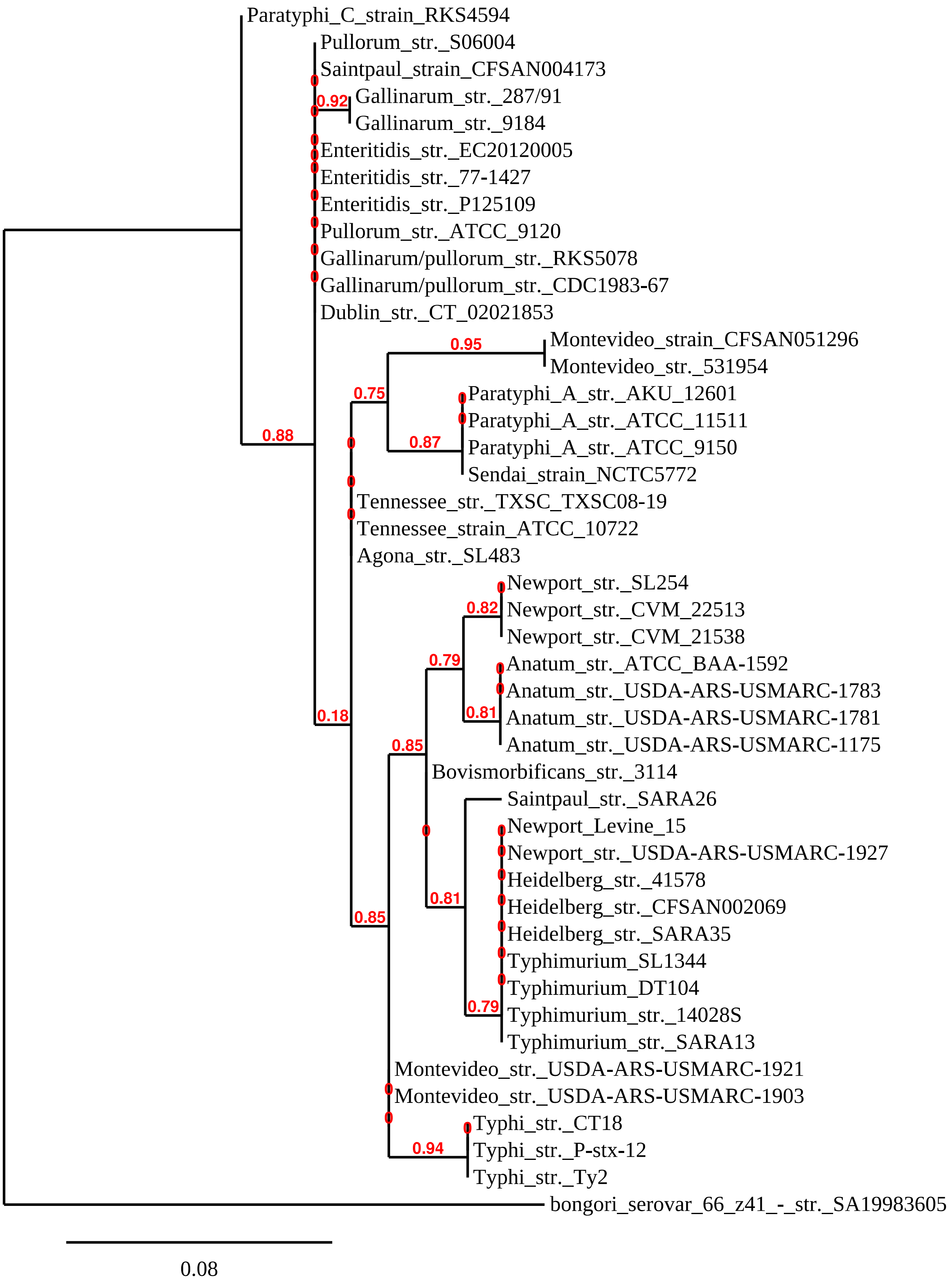

Paratyphi_C_strain_RKS4594
Pullorum_str._S06004
Saintpaul_strain_CFSAN004173
0.92
Gallinarum_str._287/91
Gallinarum_str._9184
Enteritidis_str._EC20120005
Enteritidis_str._77-1427
Enteritidis_str._P125109
Pullorum_str._ATCC_9120
Gallinarum/pullorum_str._RKS5078
Gallinarum/pullorum_str._CDC1983-67
Dublin_str._CT_02021853
0.95
Montevideo_strain_CFSAN051296
Montevideo_str._531954
0.75
Paratyphi_A_str._AKU_12601
Paratyphi_A_str._ATCC_11511
0.88
0.87
Paratyphi_A_str._ATCC_9150
Sendai_strain_NCTC5772
Tennessee_str._TXSC_TXSC08-19
Tennessee_strain_ATCC_10722
Agona_str._SL483
Newport_str._SL254
0.82
Newport_str._CVM_22513
Newport_str._CVM_21538
0.79
Anatum_str._ATCC_BAA-1592
Anatum_str._USDA-ARS-USMARC-1783
0.18
0.81
Anatum_str._USDA-ARS-USMARC-1781
Anatum_str._USDA-ARS-USMARC-1175
0.85
Bovismorbificans_str._3114
Saintpaul_str._SARA26
Newport_Levine_15
Newport_str._USDA-ARS-USMARC-1927
Heidelberg_str._41578
0.81
Heidelberg_str._CFSAN002069
0.85
Heidelberg_str._SARA35
Typhimurium_SL1344
Typhimurium_DT104
0.79
Typhimurium_str._14028S
Typhimurium_str._SARA13
Montevideo_str._USDA-ARS-USMARC-1921
Montevideo_str._USDA-ARS-USMARC-1903
Typhi_str._CT18
0.94
Typhi_str._P-stx-12
Typhi_str._Ty2
bongori_serovar_66_z41_-_str._SA19983605
0.08

**Figure S5**

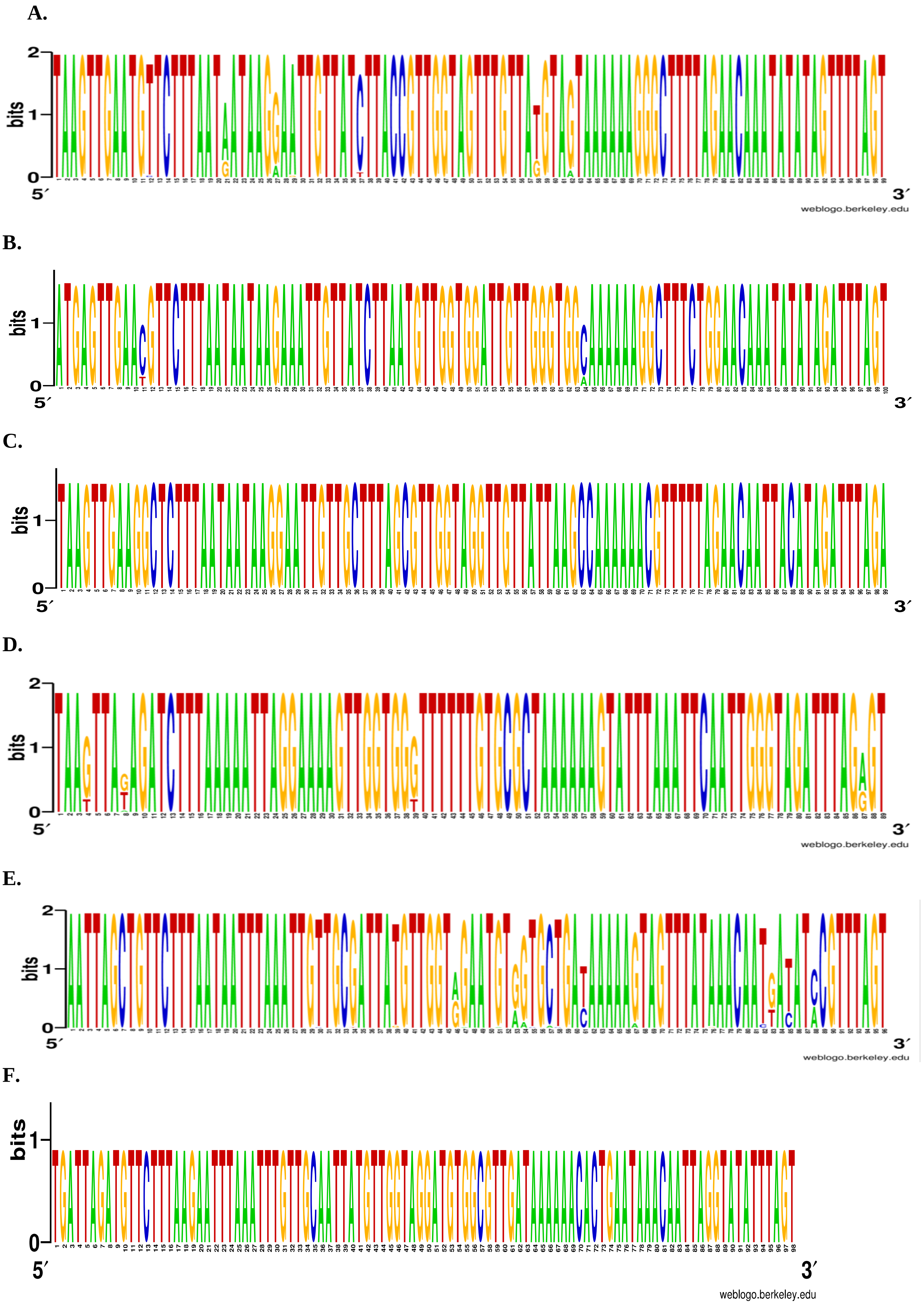

**Figure S6**

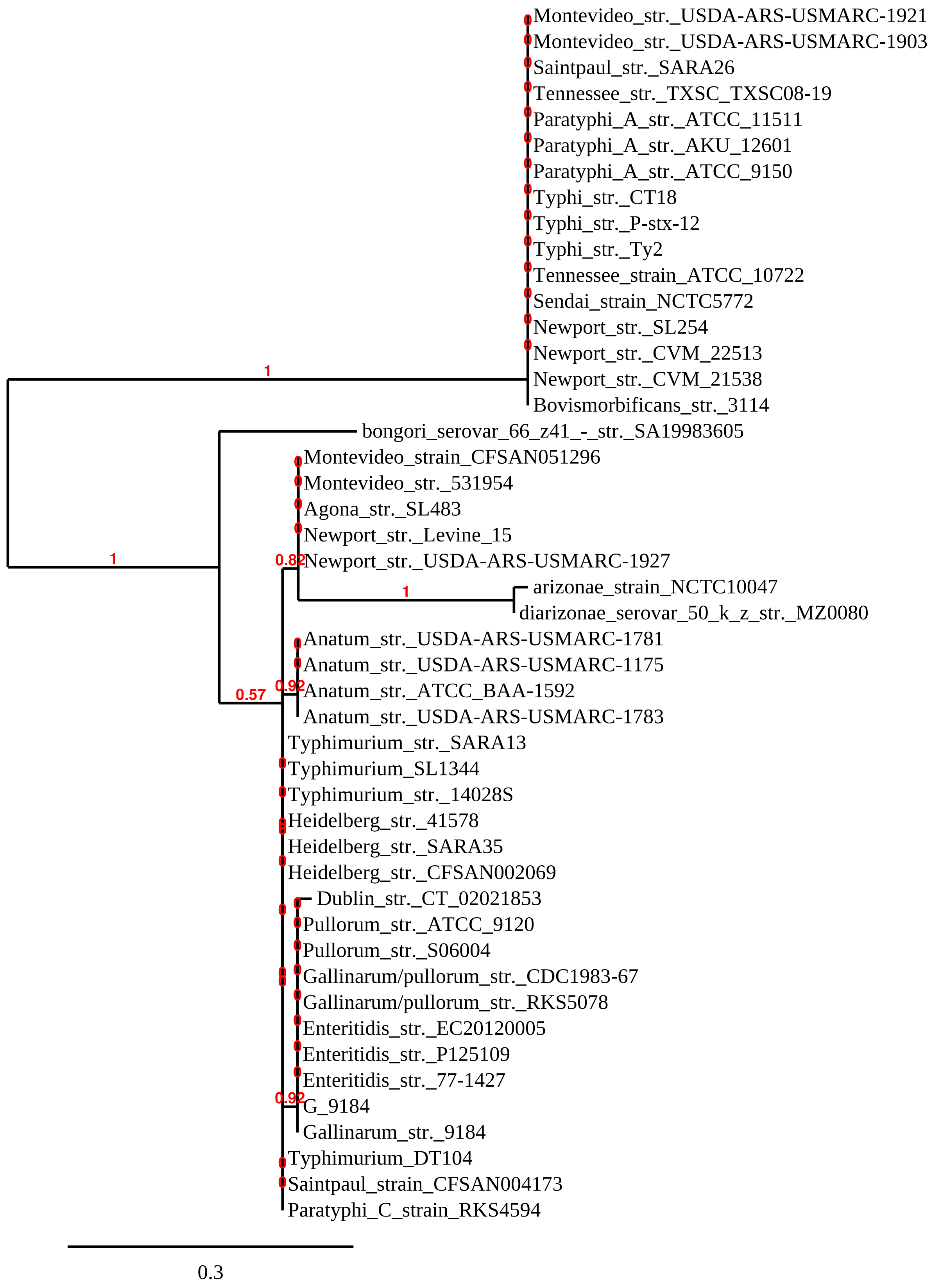
Montevideo_str._USDA-ARS-USMARC-1921
Montevideo_str._USDA-ARS-USMARC-1903
Saintpaul_str._SARA26
Tennessee_str._TXSC_TXSC08-19
Paratyphi_A_str._ATCC_11511
Paratyphi_A_str._AKU_12601
Paratyphi_A_str._ATCC_9150
Typhi_str._CT18
Typhi_str._P-stx-12
Typhi_str._Ty2
Tennessee_strain_ATCC_10722
Sendai_strain_NCTC5772
Newport_str._SL254
Newport_str._CVM_22513
Newport_str._CVM_21538
Bovismorbificans_str._3114
1
bongori_serovar_66_z41_-_str._SA19983605
Montevideo_strain_CFSAN051296
Montevideo_str._531954
Agona_str._SL483
Newport_str._Levine_15
1
0.82
Newport_str._USDA-ARS-USMARC-1927
1
arizonae_strain_NCTC10047
diarizonae_serovar_50_k_z_str._MZ0080
Anatum_str._USDA-ARS-USMARC-1781
Anatum_str._USDA-ARS-USMARC-1175
0.92
Anatum_str._ATCC_BAA-1592
0.57
Anatum_str._USDA-ARS-USMARC-1783
Typhimurium_str._SARA13
Typhimurium_SL1344
Typhimurium_str._14028S
Heidelberg_str._41578
Heidelberg_str._SARA35
Heidelberg_str._CFSAN002069
Dublin_str._CT_02021853
Pullorum_str._ATCC_9120
Pullorum_str._S06004
Gallinarum/pullorum_str._CDC1983-67
Gallinarum/pullorum_str._RKS5078
Enteritidis_str._EC20120005
Enteritidis_str._P125109
Enteritidis_str._77-1427
0.92
G_9184
Gallinarum_str._9184
Typhimurium_DT104
Saintpaul_strain_CFSAN004173
Paratyphi_C_strain_RKS4594
0.3

### Figure S7

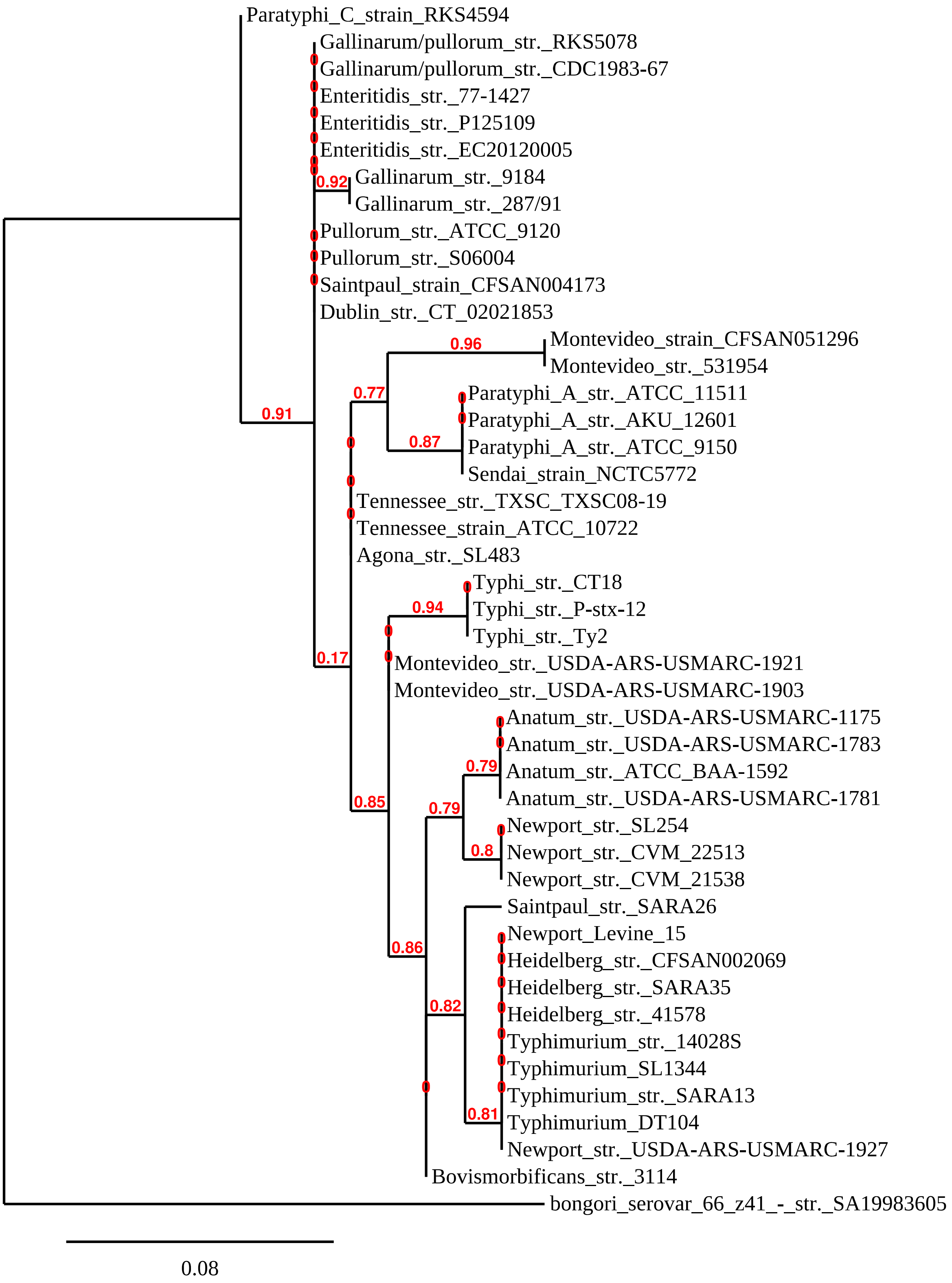
Paratyphi_C_strain_RKS4594
Gallinarum/pullorum_str._RKS5078
Gallinarum/pullorum_str._CDC1983-67
Enteritidis_str._77-1427
Enteritidis_str._P125109
Enteritidis_str._EC20120005
0.92
Gallinarum_str._9184
Gallinarum_str._287/91
Pullorum_str._ATCC_9120
Pullorum_str._S06004
Saintpaul_strain_CFSAN004173
Dublin_str._CT_02021853
0.96
Montevideo_strain_CFSAN051296
Montevideo_str._531954
0.77
Paratyphi_A_str._ATCC_11511
Paratyphi_A_str._AKU_12601
0.91
0.87
Paratyphi_A_str._ATCC_9150
Sendai_strain_NCTC5772
Tennessee_str._TXSC_TXSC08-19
Tennessee_strain_ATCC_10722
Agona_str._SL483
Typhi_str._CT18
0.94
Typhi_str._P-stx-12
Typhi_str._Ty2
0.17
Montevideo_str._USDA-ARS-USMARC-1921
Montevideo_str._USDA-ARS-USMARC-1903
Anatum_str._USDA-ARS-USMARC-1175
Anatum_str._USDA-ARS-USMARC-1783
0.79
Anatum_str._ATCC_BAA-1592
Anatum_str._USDA-ARS-USMARC-1781
0.85
0.79
Newport_str._SL254
0.8
Newport_str._CVM_22513
Newport_str._CVM_21538
Saintpaul_str._SARA26
Newport_Levine_15
0.86
Heidelberg_str._CFSAN002069
Heidelberg_str._SARA35
0.82
Heidelberg_str._41578
Typhimurium_str._14028S
Typhimurium_SL1344
Typhimurium_str._SARA13
0.81
Typhimurium_DT104
Newport_str._USDA-ARS-USMARC-1927
Bovismorbificans_str._3114
bongori_serovar_66_z41_-_str._SA19983605
0.08

**Figure 8**

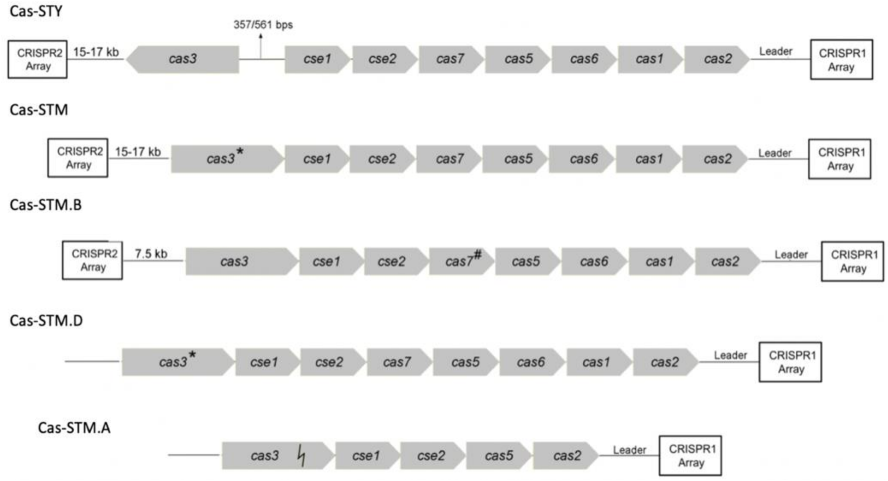


**Figure 9**

| | Cas-STM | Cas-STY | Cas-STM.A | Cas-STM.D | Cas-STM.B |
|---|---|---|---|---|---|
| Cas-STM | | 28.6 | 87 | 87 | 85 |
| Cas-STY | | | 23.5 | 24 | 29 |
| Cas-STM.A | | | | 99 | 88 |
| Cas-STM.D | | | | | 88.2 |
| Cas-STM.B | | | | | |

**Figure S10**

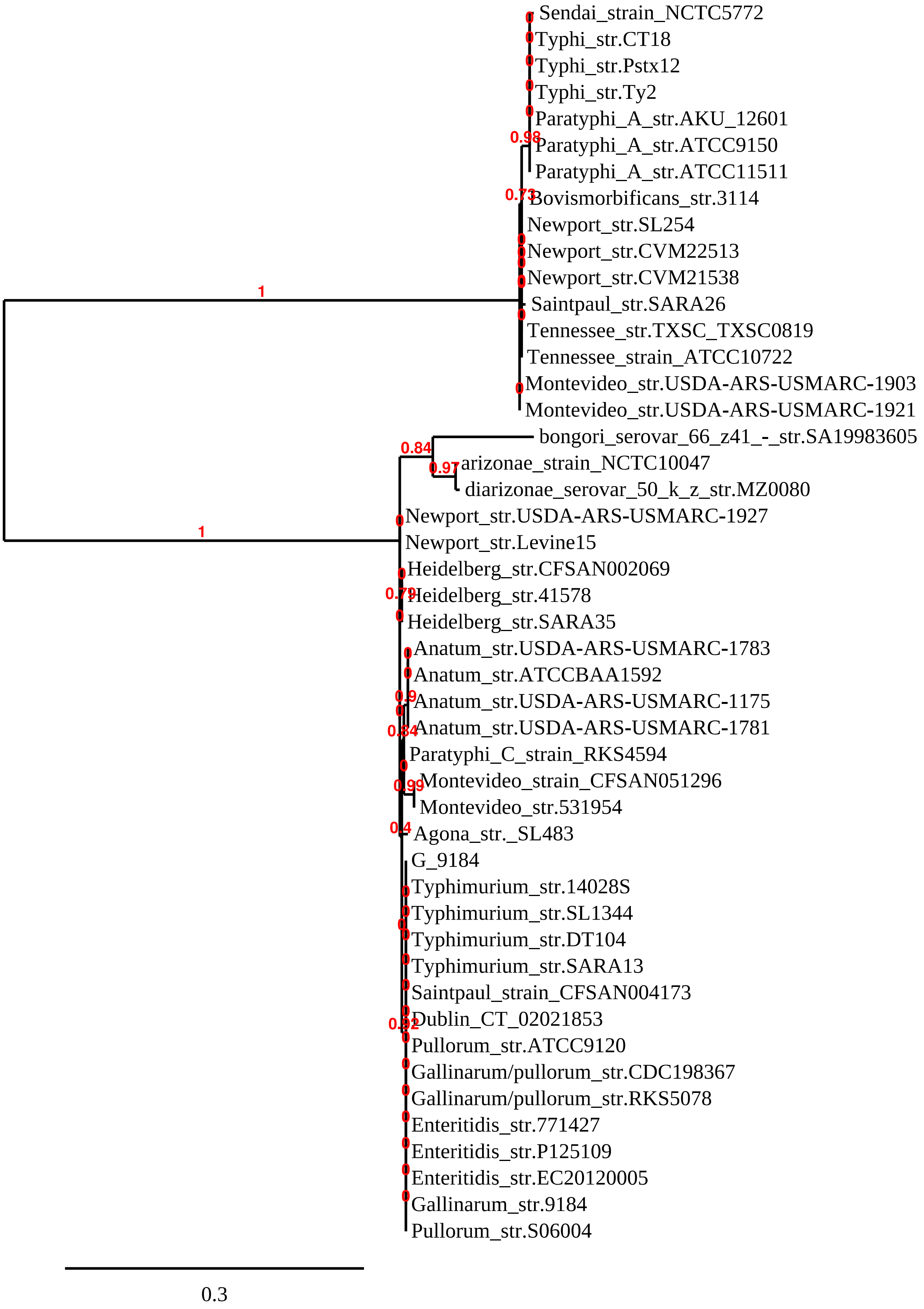

Sendai_strain_NCTC5772
Typhi_str.CT18
Typhi_str.Pstx12
Typhi_str.Ty2
Paratyphi_A_str.AKU_12601
Paratyphi_A_str.ATCC9150
Paratyphi_A_str.ATCC11511
Bovismorbificans_str.3114
Newport_str.SL254
Newport_str.CVM22513
Newport_str.CVM21538
Saintpaul_str.SARA26
Tennessee_str.TXSC_TXSC0819
Tennessee_strain_ATCC10722
Montevideo_str.USDA-ARS-USMARC-1903
Montevideo_str.USDA-ARS-USMARC-1921
bongori_serovar_66_z41_-_str.SA19983605
arizonae_strain_NCTC10047
diarizonae_serovar_50_k_z_str.MZ0080
Newport_str.USDA-ARS-USMARC-1927
Newport_str.Levine15
Heidelberg_str.CFSAN002069
Heidelberg_str.41578
Heidelberg_str.SARA35
Anatum_str.USDA-ARS-USMARC-1783
Anatum_str.ATCCBAA1592
Anatum_str.USDA-ARS-USMARC-1175
Anatum_str.USDA-ARS-USMARC-1781
Paratyphi_C_strain_RKS4594
Montevideo_strain_CFSAN051296
Montevideo_str.531954
Agona_str._SL483
G_9184
Typhimurium_str.14028S
Typhimurium_str.SL1344
Typhimurium_str.DT104
Typhimurium_str.SARA13
Saintpaul_strain_CFSAN004173
Dublin_CT_02021853
Pullorum_str.ATCC9120
Gallinarum/pullorum_str.CDC198367
Gallinarum/pullorum_str.RKS5078
Enteritidis_str.771427
Enteritidis_str.P125109
Enteritidis_str.EC20120005
Gallinarum_str.9184
Pullorum_str.S06004
1
1
0.98
0.73
0.84
0.97
0.79
0.9
0.84
0.99
0.4
0.92
0.3

### Figure S11

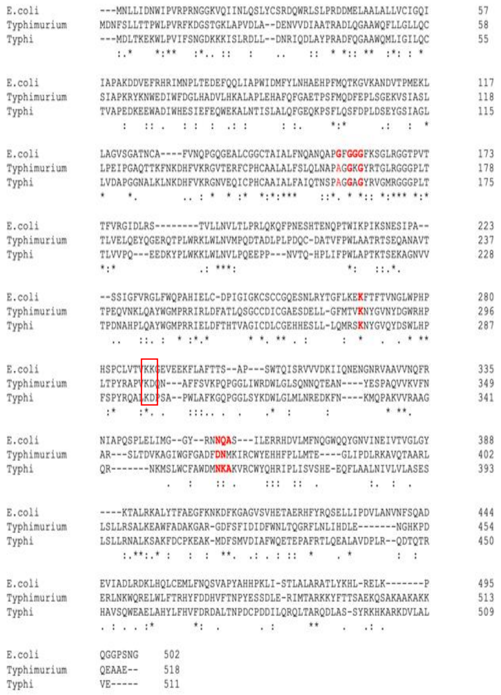

### Figure S12

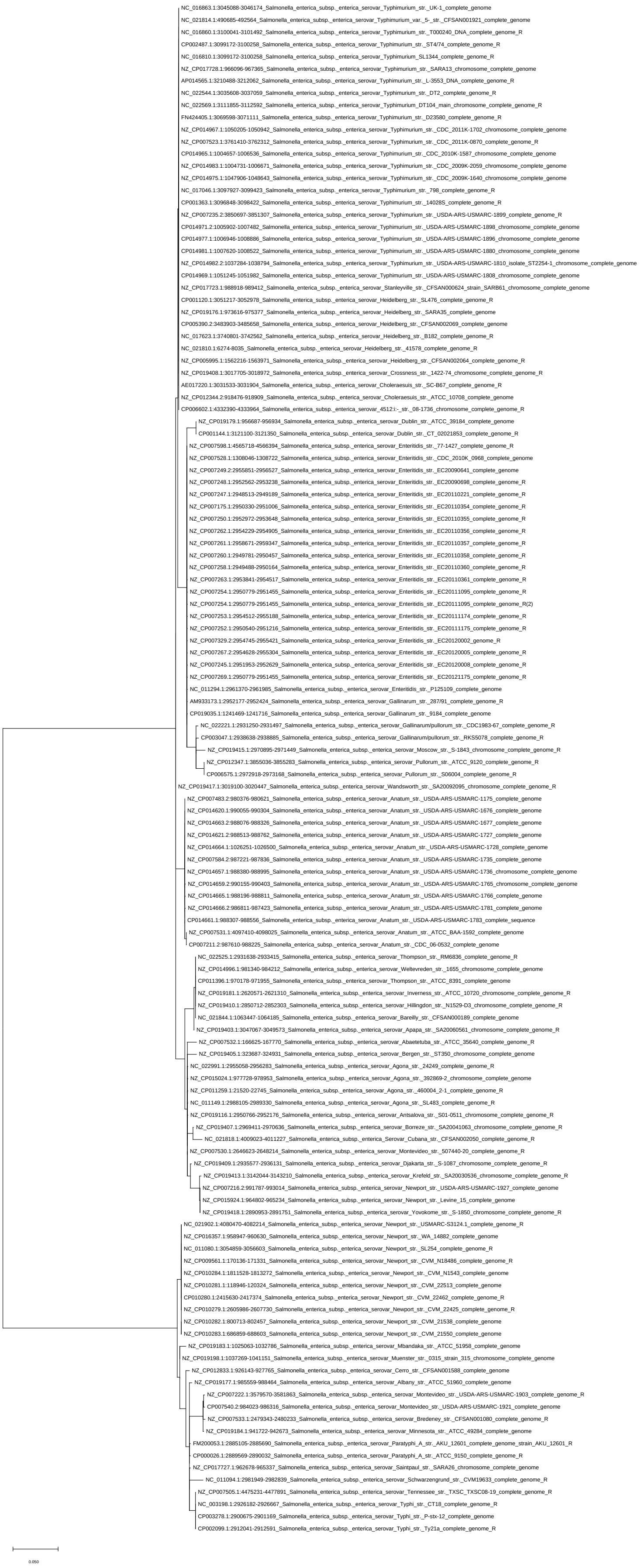
NC_016863.1:3045088-3046174_Salmonella_enterica_subsp._enterica_serovar_Typhimurium_str._UK-1_complete_genome
NC_021814.1:490685-492564_Salmonella_enterica_subsp._enterica_serovar_Typhimurium_var._5-_str._CFSAN001921_complete_genome
NC_016860.1:3100041-3101492_Salmonella_enterica_subsp._enterica_serovar_Typhimurium_str._T000240_DNA_complete_genome_R
CP002487.1:3099172-3100258_Salmonella_enterica_subsp._enterica_serovar_Typhimurium_str._ST4/74_complete_genome_R
NC_016810.1:3099172-3100258_Salmonella_enterica_subsp._enterica_serovar_Typhimurium_SL1344_complete_genome_R
NZ_CP017728.1:966096-967365_Salmonella_enterica_subsp._enterica_serovar_Typhimurium_str._SARA13_chromosome_complete_genome
AP014565.1:3210488-3212062_Salmonella_enterica_subsp._enterica_serovar_Typhimurium_str._L-3553_DNA_complete_genome_R
NC_022544.1:3035608-3037059_Salmonella_enterica_subsp._enterica_serovar_Typhimurium_str._DT2_complete_genome_R
NC_022569.1:3111855-3112592_Salmonella_enterica_subsp._enterica_serovar_Typhimurium_DT104_main_chromosome_complete_genome_R
FN424405.1:3069598-3071111_Salmonella_enterica_subsp._enterica_serovar_Typhimurium_str._D23580_complete_genome_R
NZ_CP014967.1:1050205-1050942_Salmonella_enterica_subsp._enterica_serovar_Typhimurium_str._CDC_2011K-1702_chromosome_complete_genome
NZ_CP007523.1:3761410-3762312_Salmonella_enterica_subsp._enterica_serovar_Typhimurium_str._CDC_2011K-0870_complete_genome_R
CP014965.1:1004657-1006536_Salmonella_enterica_subsp._enterica_serovar_Typhimurium_str._CDC_2010K-1587_chromosome_complete_genome
NZ_CP014983.1:1004731-1006671_Salmonella_enterica_subsp._enterica_serovar_Typhimurium_str._CDC_2009K-2059_chromosome_complete_genome
NZ_CP014975.1:1047906-1048643_Salmonella_enterica_subsp._enterica_serovar_Typhimurium_str._CDC_2009K-1640_chromosome_complete_genome
NC_017046.1:3097927-3099423_Salmonella_enterica_subsp._enterica_serovar_Typhimurium_str._798_complete_genome_R
CP001363.1:3096848-3098422_Salmonella_enterica_subsp._enterica_serovar_Typhimurium_str._14028S_complete_genome_R
NZ_CP007235.2:3850697-3851307_Salmonella_enterica_subsp._enterica_serovar_Typhimurium_str._USDA-ARS-USMARC-1899_complete_genome_R
CP014971.2:1005902-1007482_Salmonella_enterica_subsp._enterica_serovar_Typhimurium_str._USDA-ARS-USMARC-1898_chromosome_complete_genome
CP014977.1:1006946-1008886_Salmonella_enterica_subsp._enterica_serovar_Typhimurium_str._USDA-ARS-USMARC-1896_chromosome_complete_genome
CP014981.1:1007620-1008522_Salmonella_enterica_subsp._enterica_serovar_Typhimurium_str._USDA-ARS-USMARC-1880_chromosome_complete_genome
NZ_CP014982.2:1037284-1038794_Salmonella_enterica_subsp._enterica_serovar_Typhimurium_str._USDA-ARS-USMARC-1810_isolate_ST2254-1_chromosome_complete_genome
CP014969.1:1051245-1051982_Salmonella_enterica_subsp._enterica_serovar_Typhimurium_str._USDA-ARS-USMARC-1808_chromosome_complete_genome
NZ_CP017723.1:988918-989412_Salmonella_enterica_subsp._enterica_serovar_Stanleyville_str._CFSAN000624_strain_SARB61_chromosome_complete_genome
CP001120.1:3051217-3052978_Salmonella_enterica_subsp._enterica_serovar_Heidelberg_str._SL476_complete_genome_R
NZ_CP019176.1:973616-975377_Salmonella_enterica_subsp._enterica_serovar_Heidelberg_str._SARA35_complete_genome
CP005390.2:3483903-3485658_Salmonella_enterica_subsp._enterica_serovar_Heidelberg_str._CFSAN002069_complete_genome
NC_017623.1:3740801-3742562_Salmonella_enterica_subsp._enterica_serovar_Heidelberg_str._B182_complete_genome_R
NC_021810.1:6274-8035_Salmonella_enterica_subsp._enterica_serovar_Heidelberg_str._41578_complete_genome_R
NZ_CP005995.1:1562216-1563971_Salmonella_enterica_subsp._enterica_serovar_Heidelberg_str._CFSAN002064_complete_genome_R
NZ_CP019408.1:3017705-3018972_Salmonella_enterica_subsp._enterica_serovar_Crossness_str._1422-74_chromosome_complete_genome_R
AE017220.1:3031533-3031904_Salmonella_enterica_subsp._enterica_serovar_Choleraesuis_str._SC-B67_complete_genome_R
NZ_CP012344.2:918476-918909_Salmonella_enterica_subsp._enterica_serovar_Choleraesuis_str._ATCC_10708_complete_genome
CP006602.1:4332390-4333964_Salmonella_enterica_subsp._enterica_serovar_4512:i:-_str._08-1736_chromosome_complete_genome_R
NZ_CP019179.1:956687-956934_Salmonella_enterica_subsp._enterica_serovar_Dublin_str._ATCC_39184_complete_genome
CP001144.1:3121100-3121350_Salmonella_enterica_subsp._enterica_serovar_Dublin_str._CT_02021853_complete_genome_R
NZ_CP007598.1:4565718-4566394_Salmonella_enterica_subsp._enterica_serovar_Enteritidis_str._77-1427_complete_genome_R
NZ_CP007528.1:1308046-1308722_Salmonella_enterica_subsp._enterica_serovar_Enteritidis_str._CDC_2010K_0968_complete_genome
NZ_CP007249.2:2955851-2956527_Salmonella_enterica_subsp._enterica_serovar_Enteritidis_str._EC20090641_complete_genome
NZ_CP007248.1:2952562-2953238_Salmonella_enterica_subsp._enterica_serovar_Enteritidis_str._EC20090698_complete_genome_R
NZ_CP007247.1:2948513-2949189_Salmonella_enterica_subsp._enterica_serovar_Enteritidis_str._EC20110221_complete_genome_R
NZ_CP007175.1:2950330-2951006_Salmonella_enterica_subsp._enterica_serovar_Enteritidis_str._EC20110354_complete_genome_R
NZ_CP007250.1:2952972-2953648_Salmonella_enterica_subsp._enterica_serovar_Enteritidis_str._EC20110355_complete_genome_R
NZ_CP007262.1:2954229-2954905_Salmonella_enterica_subsp._enterica_serovar_Enteritidis_str._EC20110356_complete_genome_R
NZ_CP007261.1:2958671-2959347_Salmonella_enterica_subsp._enterica_serovar_Enteritidis_str._EC20110357_complete_genome_R
NZ_CP007260.1:2949781-2950457_Salmonella_enterica_subsp._enterica_serovar_Enteritidis_str._EC20110358_complete_genome_R
NZ_CP007258.1:2949488-2950164_Salmonella_enterica_subsp._enterica_serovar_Enteritidis_str._EC20110360_complete_genome_R
NZ_CP007263.1:2953841-2954517_Salmonella_enterica_subsp._enterica_serovar_Enteritidis_str._EC20110361_complete_genome_R
NZ_CP007254.1:2950779-2951455_Salmonella_enterica_subsp._enterica_serovar_Enteritidis_str._EC20111095_complete_genome_R
NZ_CP007254.1:2950779-2951455_Salmonella_enterica_subsp._enterica_serovar_Enteritidis_str._EC20111095_complete_genome_R(2)
NZ_CP007253.1:2954512-2955188_Salmonella_enterica_subsp._enterica_serovar_Enteritidis_str._EC20111174_complete_genome_R
NZ_CP007252.1:2950540-2951216_Salmonella_enterica_subsp._enterica_serovar_Enteritidis_str._EC20111175_complete_genome_R
NZ_CP007329.2:2954745-2955421_Salmonella_enterica_subsp._enterica_serovar_Enteritidis_str._EC20120002_genome_R
NZ_CP007267.2:2954628-2955304_Salmonella_enterica_subsp._enterica_serovar_Enteritidis_str._EC20120005_complete_genome_R
NZ_CP007245.1:2951953-2952629_Salmonella_enterica_subsp._enterica_serovar_Enteritidis_str._EC20120008_complete_genome_R
NZ_CP007269.1:2950779-2951455_Salmonella_enterica_subsp._enterica_serovar_Enteritidis_str._EC20121175_complete_genome_R
NC_011294.1:2961370-2961985_Salmonella_enterica_subsp._enterica_serovar_Enteritidis_str._P125109_complete_genome
AM933173.1:2952177-2952424_Salmonella_enterica_subsp._enterica_serovar_Gallinarum_str._287/91_complete_genome_R
CP019035.1:1241469-1241716_Salmonella_enterica_subsp._enterica_serovar_Gallinarum_str._9184_complete_genome
NC_022221.1:2931250-2931497_Salmonella_enterica_subsp._enterica_serovar_Gallinarum/pullorum_str._CDC1983-67_complete_genome_R
CP003047.1:2938638-2938885_Salmonella_enterica_subsp._enterica_serovar_Gallinarum/pullorum_str._RKS5078_complete_genome_R
NZ_CP019415.1:2970895-2971449_Salmonella_enterica_subsp._enterica_serovar_Moscow_str._S-1843_chromosome_complete_genome_R
NZ_CP012347.1:3855036-3855283_Salmonella_enterica_subsp._enterica_serovar_Pullorum_str._ATCC_9120_complete_genome_R
CP006575.1:2972918-2973168_Salmonella_enterica_subsp._enterica_serovar_Pullorum_str._S06004_complete_genome_R
NZ_CP019417.1:3019100-3020447_Salmonella_enterica_subsp._enterica_serovar_Wandsworth_str._SA20092095_chromosome_complete_genome_R
NZ_CP007483.2:980376-980621_Salmonella_enterica_subsp._enterica_serovar_Anatum_str._USDA-ARS-USMARC-1175_complete_genome
NZ_CP014620.1:990055-990304_Salmonella_enterica_subsp._enterica_serovar_Anatum_str._USDA-ARS-USMARC-1676_complete_genome
NZ_CP014663.2:988076-988326_Salmonella_enterica_subsp._enterica_serovar_Anatum_str._USDA-ARS-USMARC-1677_complete_genome
NZ_CP014621.2:988513-988762_Salmonella_enterica_subsp._enterica_serovar_Anatum_str._USDA-ARS-USMARC-1727_complete_genome
NZ_CP014664.1:1026251-1026500_Salmonella_enterica_subsp._enterica_serovar_Anatum_str._USDA-ARS-USMARC-1728_complete_genome
NZ_CP007584.2:987221-987836_Salmonella_enterica_subsp._enterica_serovar_Anatum_str._USDA-ARS-USMARC-1735_complete_genome
NZ_CP014657.1:988380-988995_Salmonella_enterica_subsp._enterica_serovar_Anatum_str._USDA-ARS-USMARC-1736_chromosome_complete_genome
NZ_CP014659.2:990155-990403_Salmonella_enterica_subsp._enterica_serovar_Anatum_str._USDA-ARS-USMARC-1765_chromosome_complete_genome
NZ_CP014665.1:988196-988811_Salmonella_enterica_subsp._enterica_serovar_Anatum_str._USDA-ARS-USMARC-1766_complete_genome
NZ_CP014666.2:986811-987423_Salmonella_enterica_subsp._enterica_serovar_Anatum_str._USDA-ARS-USMARC-1781_complete_genome
CP014661.1:988307-988556_Salmonella_enterica_subsp._enterica_serovar_Anatum_str._USDA-ARS-USMARC-1783_complete_sequence
NZ_CP007531.1:4097410-4098025_Salmonella_enterica_subsp._enterica_serovar_Anatum_str._ATCC_BAA-1592_complete_genome
CP007211.2:987610-988225_Salmonella_enterica_subsp._enterica_serovar_Anatum_str._CDC_06-0532_complete_genome
NC_022525.1:2931638-2933415_Salmonella_enterica_subsp._enterica_serovar_Thompson_str._RM6836_complete_genome_R
NZ_CP014996.1:981340-984212_Salmonella_enterica_subsp._enterica_serovar_Weltevreden_str._1655_chromosome_complete_genome
CP011396.1:970178-971955_Salmonella_enterica_subsp._enterica_serovar_Thompson_str._ATCC_8391_complete_genome
NZ_CP019181.1:2620571-2621310_Salmonella_enterica_subsp._enterica_serovar_Inverness_str._ATCC_10720_chromosome_complete_genome_R
NZ_CP019410.1:2850712-2852303_Salmonella_enterica_subsp._enterica_serovar_Hillingdon_str._N1529-D3_chromosome_complete_genome_R
NC_021844.1:1063447-1064185_Salmonella_enterica_subsp._enterica_serovar_Bareilly_str._CFSAN000189_complete_genome
NZ_CP019403.1:3047067-3049573_Salmonella_enterica_subsp._enterica_serovar_Apapa_str._SA20060561_chromosome_complete_genome_R
NZ_CP007532.1:166625-167770_Salmonella_enterica_subsp._enterica_serovar_Abaetetuba_str._ATCC_35640_complete_genome_R
NZ_CP019405.1:323687-324931_Salmonella_enterica_subsp._enterica_serovar_Bergen_str._ST350_chromosome_complete_genome
NC_022991.1:2955058-2956283_Salmonella_enterica_subsp._enterica_serovar_Agona_str._24249_complete_genome_R
NZ_CP015024.1:977728-978953_Salmonella_enterica_subsp._enterica_serovar_Agona_str._392869-2_chromosome_complete_genome
NZ_CP011259.1:21520-22745_Salmonella_enterica_subsp._enterica_serovar_Agona_str._460004_2-1_complete_genome_R
NC_011149.1:2988105-2989330_Salmonella_enterica_subsp._enterica_serovar_Agona_str._SL483_complete_genome_R
NZ_CP019116.1:2950766-2952176_Salmonella_enterica_subsp._enterica_serovar_Antsalova_str._S01-0511_chromosome_complete_genome_R
NZ_CP019407.1:2969411-2970636_Salmonella_enterica_subsp._enterica_serovar_Borreze_str._SA20041063_chromosome_complete_genome_R
NC_021818.1:4009023-4011227_Salmonella_enterica_subsp._enterica_Serovar_Cubana_str._CFSAN002050_complete_genome_R
NZ_CP007530.1:2646623-2648214_Salmonella_enterica_subsp._enterica_serovar_Montevideo_str._507440-20_complete_genome_R
NZ_CP019409.1:2935577-2936131_Salmonella_enterica_subsp._enterica_serovar_Djakarta_str._S-1087_chromosome_complete_genome_R
NZ_CP019413.1:3142044-3143210_Salmonella_enterica_subsp._enterica_serovar_Krefeld_str._SA20030536_chromosome_complete_genome_R
NZ_CP007216.2:991787-993014_Salmonella_enterica_subsp._enterica_serovar_Newport_str._USDA-ARS-USMARC-1927_complete_genome
NZ_CP015924.1:964802-965234_Salmonella_enterica_subsp._enterica_serovar_Newport_str._Levine_15_complete_genome
NZ_CP019418.1:2890953-2891751_Salmonella_enterica_subsp._enterica_serovar_Yovokome_str._S-1850_chromosome_complete_genome_R
NC_021902.1:4080470-4082214_Salmonella_enterica_subsp._enterica_serovar_Newport_str._USMARC-S3124.1_complete_genome_R
NZ_CP016357.1:958947-960630_Salmonella_enterica_subsp._enterica_serovar_Newport_str._WA_14882_complete_genome
NC_011080.1:3054859-3056603_Salmonella_enterica_subsp._enterica_serovar_Newport_str._SL254_complete_genome_R
NZ_CP009561.1:170136-171331_Salmonella_enterica_subsp._enterica_serovar_Newport_str._CVM_N18486_complete_genome_R
NZ_CP010284.1:1811528-1813272_Salmonella_enterica_subsp._enterica_serovar_Newport_str._CVM_N1543_complete_genome
NZ_CP010281.1:118946-120324_Salmonella_enterica_subsp._enterica_serovar_Newport_str._CVM_22513_complete_genome
CP010280.1:2415630-2417374_Salmonella_enterica_subsp._enterica_serovar_Newport_str._CVM_22462_complete_genome_R
NZ_CP010279.1:2605986-2607730_Salmonella_enterica_subsp._enterica_serovar_Newport_str._CVM_22425_complete_genome_R
NZ_CP010282.1:800713-802457_Salmonella_enterica_subsp._enterica_serovar_Newport_str._CVM_21538_complete_genome
NZ_CP010283.1:686859-688603_Salmonella_enterica_subsp._enterica_serovar_Newport_str._CVM_21550_complete_genome
NZ_CP019183.1:1025063-1032786_Salmonella_enterica_subsp._enterica_serovar_Mbandaka_str._ATCC_51958_complete_genome
NZ_CP019198.1:1037269-1041151_Salmonella_enterica_subsp._enterica_serovar_Muenster_str._0315_strain_315_chromosome_complete_genome
NZ_CP012833.1:926143-927765_Salmonella_enterica_subsp._enterica_serovar_Cerro_str._CFSAN001588_complete_genome
NZ_CP019177.1:985559-988464_Salmonella_enterica_subsp._enterica_serovar_Albany_str._ATCC_51960_complete_genome
NZ_CP007222.1:3579570-3581863_Salmonella_enterica_subsp._enterica_serovar_Montevideo_str._USDA-ARS-USMARC-1903_complete_genome_R
CP007540.2:984023-986316_Salmonella_enterica_subsp._enterica_serovar_Montevideo_str._USDA-ARS-USMARC-1921_complete_genome
NZ_CP007533.1:2479343-2480233_Salmonella_enterica_subsp._enterica_serovar_Bredeney_str._CFSAN001080_complete_genome_R
NZ_CP019184.1:941722-942673_Salmonella_enterica_subsp._enterica_serovar_Minnesota_str._ATCC_49284_complete_genome
FM200053.1:2885105-2885690_Salmonella_enterica_subsp._enterica_serovar_Paratyphi_A_str._AKU_12601_complete_genome_strain_AKU_12601_R
CP000026.1:2889569-2890032_Salmonella_enterica_subsp._enterica_serovar_Paratyphi_A_str._ATCC_9150_complete_genome_R
NZ_CP017727.1:962678-965337_Salmonella_enterica_subsp._enterica_serovar_Saintpaul_str._SARA26_chromosome_complete_genome
NC_011094.1:2981949-2982839_Salmonella_enterica_subsp._enterica_serovar_Schwarzengrund_str._CVM19633_complete_genome_R
NZ_CP007505.1:4475231-4477891_Salmonella_enterica_subsp._enterica_serovar_Tennessee_str._TXSC_TXSC08-19_complete_genome_R
NC_003198.1:2926182-2926667_Salmonella_enterica_subsp._enterica_serovar_Typhi_str._CT18_complete_genome_R
CP003278.1:2900675-2901169_Salmonella_enterica_subsp._enterica_serovar_Typhi_str._P-stx-12_complete_genome
CP002099.1:2912041-2912591_Salmonella_enterica_subsp._enterica_serovar_Typhi_str._Ty21a_complete_genome_R
0.050

**Supplementary Methodology**

***CRISPR loci data collection in correct orientation***: Our study comprises 131 strains belonging to two species, *S. bongori*, and *S. enterica*, including 22 serovars and three subspecies (supplementary table S1). These samples were primitively isolated from multiple sources, including primates, poultry, swine, cattle, food specimens, and natural environment (GenBank database). The complete genome sequences for all these annotated strains were obtained from the GenBank database. Only experimentally validated sequences were considered to ensure the legitimacy of the data being used. The CRISPR loci were identified in two steps - the annotation and orientation of the CRISPR array were retrieved from the online database of CRISPR- Cas++ (Couvin et al. 2018). The upstream and downstream regions of these arrays were aligned with the leader sequences previously reported by (Shariat et al. 2015) to know the correct sequence of the CRISPR array. The arrays were then classified as CRISPR1 and CRISPR2 after verifying the leader sequence and its position with respect to the *cas* operon (Shariat et al. 2015).

Most strains of *S. enterica* subsp. *enterica* had both, the CRISPR1 and CRISPR2 arrays. However, all the analyzed strains of *S. enterica* subsp. *enterica* serovar Heidelberg, a few strains of serovar Typhimurium, and one strain of serovar Tennessee are reported to harbor more than two CRISPR arrays (Couvin et al. 2018). Instead, our analysis confirmed that the CRISPR1 array of serovars Typhimurium and Heidelberg were divided into two parts by a stretch of 74 nucleotides consisting of two truncated spacers and a direct repeat (DR) (supplementary fig. S1A). The two parts of the CRISPR1 array taken together in concatenation aligned well with the intact CRISPR1 array of other strains of serovar Typhimurium. Similarly, the CRISPR1 array of serovar Tennessee strain (str.) CFSAN070645 was divided into three parts (containing 19, 24, and 16 spacers) and the CRISPR2 into two parts (consisting 10 and 11 spacers) due to the presence of mutated DRs rendering a stretch of 91bp undetectable as a part of the CRISPR array. Therefore, we considered the concatenated forms of these CRISPR arrays as a single unit for further analysis. Our analysis also indicated the occurrence of CRISPR1 array with two spacers each, in the serovars Dublin, Gallinarum, Pullorum, and Gallinarum/Pullorum. However, neither of these CRISPR arrays were described as valid in the CRISPR-Cas++ database, and the CRISPRCasFinder software allocated 27bp long DRs and 34bp long spacer sequences. Likewise, the CRISPR2 arrays of serovar Typhi and serovar Pullorum str. S06004 identified through our analysis were not detectable by this database. The CRISPR2 array of serovar Typhi possessed only one erratic spacer and that of serovar Pullorum str. S06004 had two spacers. We considered all these strains and their respective CRISPR-Cas systems in our analysis.
